# Structure of long-range direct and indirect spinocerebellar pathways as well as local spinal circuits mediating proprioception

**DOI:** 10.1101/2020.08.17.254607

**Authors:** Iliodora V. Pop, Felipe Espinosa, Cheasequah J. Blevins, Portia C. Okafor, Osita W. Ogujiofor, Megan Goyal, Bishakha Mona, Mark A. Landy, Kevin M. Dean, Channabasavaiah B. Gurumurthy, Helen C. Lai

## Abstract

Proprioception, the sense of limb and body position, generates a map of the body that is essential for proper motor control, yet we know little about precisely how neurons in proprioceptive pathways are wired. Defining the anatomy of secondary neurons in the spinal cord that integrate and relay proprioceptive and potentially cutaneous information from the periphery to the cerebellum is fundamental to understanding how proprioceptive circuits function. Here, we use genetic tools in both male and female mice to define the unique anatomical trajectories of long-range direct and indirect spinocerebellar pathways as well as local intersegmental spinal circuits. We find that Clarke’s column (CC) neurons, a major contributor to the direct spinocerebellar pathway, has mossy fiber terminals that diversify extensively in the cerebellar cortex with axons terminating bilaterally, but with no significant axon collaterals within the spinal cord, medulla, or cerebellar nuclei. By contrast, we find that two of the indirect pathways, the spino-lateral reticular nucleus (spino-LRt) and spino-olivary pathways, are in part, derived from cervical *Atoh1*-lineage neurons, while thoracolumbar *Atoh1*-lineage neurons project mostly locally within the spinal cord. Notably, while cervical and thoracolumbar *Atoh1*-lineage neurons connect locally with motor neurons, no CC to motor neuron connections were detected. Silencing of caudal *Atoh1*- lineage neurons results in a subtle motor impairment consistent with a defect in local proprioceptive circuitry. Altogether, we define anatomical differences between long-range direct, indirect, and local proprioceptive subcircuits that likely mediate different components of proprioceptive-motor behaviors.

**Significance Statement:** We define the anatomy of long-range direct and indirect spinocerebellar pathways as well as local spinal proprioceptive circuits. We observe that mossy fiber axon terminals of Clarke’s column (CC) neurons diversify proprioceptive information across granule cells in multiple lobules on both ipsilateral and contralateral sides sending no significant collaterals within the spinal cord, medulla, or cerebellar nuclei. Strikingly, we find that cervical spinal cord *Atoh1*-lineage neurons form mainly the indirect spino- lateral reticular nucleus and spino-olivary tracts and thoracolumbar *Atoh1*-lineage neurons project locally within the spinal cord while only a few *Atoh1*-lineage neurons form a direct spinocerebellar tract. Altogether, we define the development, anatomical projections, and some behavioral consequences of silencing spinal proprioceptive pathways.

## Introduction

Proprioception, the sense of limb and body position, is critical for generating an online state body map (Sherrington, 1906; Tuthill and Azim, 2018). When proprioception is lost, gross trajectories are maintained, but coordinated limb movement is impaired (Gordon et al., 1995; Abelew et al., 2000; Windhorst, 2007; Akay et al., 2014). Muscle and tendon information detected by proprioceptive sensory neurons are integrated by secondary neurons in the spinal cord and relayed to the cerebellum through both direct and indirect spinocerebellar pathways (Oscarsson, 1965; Bosco and Poppele, 2001; Jiang et al., 2015). In this study, we sought to define the precise anatomy of the proprioceptive system through the spinal cord using genetic tools in mice.

The direct spinocerebellar pathway consists of ipsilaterally-projecting dorsal and contralaterally- projecting ventral spinocerebellar tracts (DSCT and VSCT), deriving from several anatomically and molecularly distinct groups of soma in diverse laminae throughout the spinal cord where they are thought to convey ongoing locomotor activity (Matsushita and Hosoya, 1979; Sengul et al., 2015). A major contributor to the DSCT comes from Clarke’s column (CC) neurons, whose soma reside in the medial aspect of the thoracic to upper lumbar spinal cord (Oscarsson, 1965; Baek et al., 2019). While SCT axons terminate as mossy fiber (MF) terminals on granule cells (GCs) in the cerebellum (Arsenio Nunes and Sotelo, 1985; Reeber et al., 2011), the extent to which CC neurons send axon collaterals to areas within the spinal cord, medulla, or cerebellar nuclei (CN) is unclear (Ekerot and Oscarsson, 1976; Matsushita and Gao, 1997; Mogensen et al., 2017; Luo et al., 2018). Such axon collaterals would be important for integration with other ascending or descending pathways (Sillitoe et al., 2012; Beitzel et al., 2017).

Compared to the direct spinocerebellar pathways, less is known about the anatomy of the indirect spino-LRt and spino-olivary (Helweg’s tract) pathways. Spino-LRt neurons project ipsilaterally and contralaterally to the LRt in the medulla where they are involved in posture, reaching, and grasping (Alstermark and Ekerot, 2013; Jiang et al., 2015). Spino-olivary neurons reside in lamina V-VII of the spinal cord and project contralaterally to the inferior olive (IO) in the medulla (Oscarsson and Sjolund, 1977a, b; Berkley and Worden, 1978; Swenson and Castro, 1983b, a). Neurons in the IO are thought to be involved in the timing of motor commands, motor learning, and error correction (Sillitoe et al., 2012; White and Sillitoe, 2017).

Developmentally, the basic helix-loop-helix (bHLH) transcription factor-expressing progenitor domains *Atoh1* (atonal homolog 1) and the dorsal *Neurog1* (neurogenin 1) domains are reported to differentiate into neurons of the direct spinocerebellar pathway (Bermingham et al., 2001; Gowan et al., 2001; Sakai et al., 2012). The soma of *Atoh1*-lineage neurons reside medially, laterally, and ventrally to CC neurons suggesting that CC neurons do not develop from *Atoh1*-lineage neurons, but from an alternate progenitor domain (Yuengert et al., 2015). Based on the location of *Atoh1*-lineage neurons, we previously hypothesized that they make other direct spinocerebellar neurons such as the lamina V-SCTs or dorsal horn-SCTs (Matsushita and Hosoya, 1979; Edgley and Gallimore, 1988).

In this study, we sought to provide clarity to the development and anatomy of the proprioceptive system using genetic labeling strategies, whole tissue clearing and imaging, and tracing tools of CC and spinal cord *Atoh1*-lineage neurons. We find that CC neurons develop from a *Neurog1*, not *Atoh1*, progenitor population. We also find that as a population, CC axons do not collateralize considerably to any structures within the spinal cord, medulla, or CN, although they do collateralize extensively within the cerebellar cortex with some axons crossing the midline within the cerebellum. Furthermore, we find that cervical *Atoh1*-lineage neurons make the indirect spino-LRt and spino-olivary tracts rather than the direct lamina V-SCTs or dorsal horn SCTs as originally hypothesized and that thoracolumbar *Atoh1*-lineage neurons project mainly locally within the spinal cord. Altogether, we provide novel insights into the development, anatomy, and function of spinal cord proprioceptive pathways.

## Materials & Methods

### Mouse strains

The following mouse strains were used: *Gdnf^IRES2-CreERT2^* (Cebrian et al., 2014)(abbreviated *Gdnf^CreER^*, JAX #024948), *Neurog1BAC-Cre* (Quinones et al., 2010)(JAX #012859), *Atoh1^P2A-FLPo^* (Ogujiofor et al., 2021), *Atoh1^Cre^* knock-in (Yang et al., 2010), *R26^LSL-LacZ^* (Soriano, 1999)(JAX #003474), *R26^LSL-tdTom^* (Ai14)(JAX #007914)(Madisen et al., 2010), *R26^LSL-FSF-tdTom^* (Ai65)(JAX #032864)(Madisen et al., 2015), *Cdx2::FLPo* (Bourane et al., 2015), *R26^LSL-FSF-TeTx^* (Kim et al., 2009). All mice were outbred and thus, are mixed strains (including C57Bl/6J, C57Bl/6N, and ICR). *Atoh1^Cre/+^* knock-in mice crossed to *Cdx2::FLPo* and dual recombinase tdTomato reporter Ai65 mice were screened for “dysregulated” expression as previously reported (Yuengert et al., 2015). Tamoxifen (Sigma) was injected at P7 and/or P8 for the *Gdnf^CreER^* line at 0.1 mg/g mouse using 10 mg/mL Tamoxifen dissolved in sunflower oil (Sigma) with 10% ethanol. All animal experiments were approved by the Institutional Animal Care and Use Committee at UT Southwestern.

### Tissue processing

Mice are age P0 on the day of birth. Mice older than P10 were anesthetized with Avertin (2,2,2- Tribromoethanol) (0.025-0.030 mL of 0.04 M Avertin in 2-methyl-2-butanol and distilled water/g mouse) and transcardially perfused, first with 0.012% w/v Heparin/PBS and then 4% PFA/PBS. A dorsal or ventral laminectomy exposed the spinal cord to the fixative. Spinal cords were fixed for 2 hrs and the brains overnight at 4°C. Tissue was washed in PBS for at least one day and cryoprotected in 30% sucrose dissolved in deionized water. Tissue was marked with 1% Alcian Blue in 3% acetic acid on one side to keep orientation and were embedded in OCT (Tissue-Tek Optimal Cutting Temperature compound). Tissue was sectioned using a Leica CM1950 Cryostat.

### Immunohistochemistry (IHC) and confocal imaging

Cryosections (20-40 μm) were blocked with PBS/1-3% normal goat or donkey serum (Jackson labs)/0.3% Triton X-100 (Sigma) for up to 1 hr at room temperature (RT) and incubated overnight with primary antibody at 4°C. After washing 3 times with PBS, the appropriate secondary antibody (Alexa 488, 567, and/or 647, Invitrogen) was incubated for an hour at RT. Sections were rinsed 3 times in PBS, mounted with Aqua-Poly/Mount (Polysciences Inc.), and coverslipped (Fisher). The following primary antibodies and dilutions were used: 1:500 rabbit anti-dsRed (Clontech), 1:500 mouse anti-NEUN (Millipore Sigma), 1:500 chicken anti-GFP (Aves), 1:5000 guinea pig anti-VGLUT1 (Millipore Sigma), 1:1000 guinea pig anti-VGLUT2 (Millipore Sigma), 1:100 goat anti-CHAT (Millipore Sigma), 1:200 rabbit anti-PSD-95 (Invitrogen), 1:1000 rabbit anti-PV27 (Swant). Sections were referenced to the Mouse Brain Atlas (Paxinos and Franklin, 2007) and Christopher Reeves Spinal Cord Atlas (Watson et al., 2009).

Fluorescent images were taken on a Zeiss LSM710 or LSM880 confocal microscope with an optical slice of 0.5-10 μm depending on the objective used (10x air, 20x air, 40x oil, or 63x oil). Images were pseudocolored using a magenta/yellow/blue, magenta/green/blue, or magenta/yellow/cyan color scheme using Adobe Photoshop 2021 (Adobe) or Fiji (Schindelin et al., 2012). Images for quantitation of soma size were processed in Fiji. Images were thresholded and the soma manually outlined to obtain the soma area.

### In situ hybridization (ISH)

ISH was performed as per standard protocols. Detailed protocol is available upon request. Briefly, spinal cord sections (30 μm) were dried at 50^°^C for 15 min. then fixed in 4% paraformaldehyde (PFA) in DEPC-PBS for 20 min. at RT. The sections were washed in DEPC-PBS for 5 min. at RT before and after the incubation in RIPA buffer (150 mM NaCl, 1% NP-40, 0.5% Na deoxycholate, 0.1% SDS, 1 mM EDTA, 50 mM Tris pH 8.0) for 60 min. Next, the sections were postfixed in 4% PFA in DEPC-PBS for 15 min at RT. The sections were then washed in DEPC-water followed by acetylation (500 µL of acetic anhydride in 200 mL of 0.1 M RNase-free triethanolamine-HCl at pH 8.0), washed in DEPC-PBS for 5 min., and prehybridized for 2 h at 60-62°C. Sections were incubated overnight at 60-62^°^C with 1–2 ng/µL of fresh probe (*Gdnf* or *Vglut1*). A series of low and high stringency washes in 2x and 0.2x SSC as well as treatment with RNaseA and RNase T1 were performed. The sections were blocked in 10% inactivated sheep serum for 1 h followed by overnight incubation with 1:1000 anti-digoxygenin (DIG) antibody (Roche). The sections were washed in PBT and incubated with NBT/BCIP (Roche) staining solution. After the blue precipitate formed, the slides were washed in PBS and coverslipped with Aqua-Poly/Mount (Polysciences Inc.) mounting media.

The RNAscope Fluorescent Multiplex Assay (Advanced Cell Diagnostics Inc., Hayward, CA) was performed according to the manufacturer’s instructions using a *Vglut1* probe (ACDBio, 416631), *Vglut2* probe (ACDBio, 319171), or *Gdnf* probe (ACDBio, 421941). All incubation steps were performed in a HybEZ^TM^ II oven set to 40°C. The slides were then washed with 1x DPBS (Gibco, 14190) and incubated with a 2:1 1x DPBS:Protease III for 150 sec. Slides were then washed with 1x DPBS three times and incubated with the probe(s) for 2 hours. The slides were washed two times thoroughly using 1x wash buffer (ACDBio, 310091) for 2 min, then incubated with Amp 1-Fl for 30 minutes. The same process (washing then treatment) was repeated for Amp 2-Fl, Amp 3-Fl and Amp 4-Fl for 15, 30 and 15 minutes, respectively. For antibody staining of β-galactosidase, the sections were transferred to a humidified tray and blocked for 30-45 minutes in 0.25mL/slide of PBT (PBS with 0.3% Triton) containing 1% goat serum (Jackson ImmunoResearch). The sections were incubated with chicken anti-β-Galactosidase antibody (Abcam, 1:500) in PBT with 1% goat serum overnight at 4°C. The slides were then washed three times in PBS for 10 minutes and incubated at room temperature for 1 hour with goat anti-chicken Alexa Fluor 488 (Life Technologies, 1:500). Slides were washed three times in PBS for 10 minutes and coverslipped using 2 drops of Aqua-Poly/Mount (Polysciences, Inc.) as the mounting media.

### X-gal staining

Slides with spinal cord sections (30 µm) were incubated in staining solution with 5mM K3Fe(CN)6, 5mM K4Fe(CN)6 and 1 mg/mL of X-gal (Roche) until precipitate was sufficient to visualize. Sections were moved to PBS, mounted, and coverslipped.

### Viral Injections

Mice aged P7-P8 were anesthetized using isoflurane (Henry Schein) and prepared for injections into the spinal cord. The back hair was shaved and 70% ethanol and betadine (Avrio Health L.P.) applied. A midline incision was made on the dorsal surface of the spinal cord. AAV9.hSyn.DIO.eGFP.WPRE.hGH was injected into the lower thoracic spinal cord through the intervertebral space of P7 or P8 *Gdnf^Tom^* mice (100 nL total in 27.6 nL increments at 1-2 min intervals (Nanoject II, Drummond Scientific), 1.07 x 10^13^ GC/mL, Penn Vector Core). Lenti^FugE^-Cre was injected into the cervical or lower thoracic to lumbar area of P7 *Atoh1^P2A-FLPo^;Ai65* mice (total of 50.6 nL in 27.6 nL increments at 1-2 min intervals). Lenti^FugE^-Cre was pseudotyped with a fusion glycoprotein enabling efficient retrograde axonal transport (Kato et al., 2014). To generate Lenti^FugE^-Cre, *Cre* was sub-cloned into the third generation HIV-based lentivirus vector under the control of a synapsin promoter (FSW-*Cre*). FSW-*Cre* was co-transfected into HEK293 cells with three packing plasmids, pMDLg/pRRE, pRSV-Rev and pCAGGS-FuG-E to generate Lenti^FugE^- Cre, which was concentrated with ultracentrifugation to 2.0 x 10^12^ Vg/mL. The incision was closed with surgical glue (Henry Schein). Carprofen (5 mg/kg) was administered daily 3 days after surgery. Spinal cords were harvested approximately 3 weeks after injection.

### CTB and FG Injections

*Gdnf^Tom^* mice were injected with 1% (w/v) of CTB-488 (left side) and CTB-647 (right side) (Thermo Fisher Scientific). Mice were anesthetized with isoflurane and the area above and around the cerebellar region was prepared for surgery. A midline incision of 0.75 cm and a craniectomy of approximately 1 mm X 1 mm was performed. Bilateral injections at 4 sites were done at (from Bregma): rostrocaudal -5.7 and -6.2 mm and at mediolateral ± 0.35 mm. At each site, several injections in 32 nL increments were performed every 300 μm along the dorsoventral axis at coordinates: -1.8 and -1.5 mm deep for a total of 320 nL of conjugated CTB. Animals were euthanized and tissue harvested 5 days after injection.

Fluorogold (FG) was injected into the vermis of lobules I-V in the cerebella of *Gdnf^Tom^*, *Atoh1^Tom^*, or *Atoh1^Cre^;Cdx2::FLPo;Ai65* mice. Mice (1-2 months old) were injected with 4% (w/v) FG solution in saline (Fluorochrome). Mice were anesthetized with isoflurane and the area above and around the cerebellar region was prepared for surgery. A midline incision of ∼0.75 cm and a craniectomy of approximately 1 mm wide by 1.5 mm long was performed. Bilateral injections at six sites were done at (from Bregma): rostrocaudal -5.6 to -5.5, -5.9, and -6.25 to -6.3 mm and at mediolateral ± 0.2-0.4 mm. The maximum depth at each rostrocaudal site was -2.0 mm, -2.4 mm, -1.5 mm, respectively. Multiple injections were done at each site in 32 or 50.6 nL increments every 300 μm along the dorsoventral axis for a total of 270-720 nL of FG on each side. Animals were euthanized and tissue harvested 7 days after injection.

For FG injections targeting the LRt and IO, *Atoh1^Tom^* mice (7-9 weeks old) were anesthetized with isoflurane and the area above and behind the occipital region prepared for surgery. A midline incision of ∼1 cm was made. Neck muscles were detached from the occipital bone and retracted laterally to expose the foramen magnum. A needle with a rostroventral inclination of 47° was used to advance through the foramen magnum into the brainstem ∼3.7 mm from dura. 32.2 nL of 1-2% (w/v) FG solution was injected 4-5 times 25 μm apart while retracting the pipette with an interval of 30 seconds between injections for a total of 128.8-161 nL of FG. Unilateral injections were done at the following coordinates measured from the most ventral aspect of the occipital crest: rostrocaudal -0.4 mm, and mediolateral ± 0.75 mm. After the last injection, the needle was left in place for 3 minutes and then slowly extracted from the brainstem. Animals were euthanized and tissue was harvested 7 days after injection.

### Whole tissue imaging

Mouse hindbrain and spinal cords were processed following the SHIELD protocol (Park et al., 2018). Tissues were cleared with SmartClear II Pro (LifeCanvas Technologies, Cambridge, MA) for several days, mounted in a gel of 0.9% agarose in EasyIndex (LifeCanvas Technologies), and then incubated in EasyIndex for refractive index matching. Tissues were imaged at 3.6X using a SmartSPIM light sheet microscope (LifeCanvas Technologies). Spinal cords and hindbrains of three mice marking CC were used for quantitation of the MF/cell body ratio: one *Gdnf^Tom^* mouse (female, P23) and two *Gdnf^Tom^*;*Cdx::FLPo;Ai65* mice (one male, one female, P28). Mice were imaged with 1.8 μm x 1.8 μm x 2 μm sampling (X, Y, and Z, respectively). The total number of 2 μm image slices for each sample was as follows: spinal cord (1500, 3300, 2400 slices) and hindbrain (2700, 3000, 3300 slices) for the *Gdnf^Tom^*, *Gdnf^Tom^*;*Cdx::FLPo;Ai65* male, *Gdnf^Tom^*;Cdx::FLPo;Ai65 female mouse, respectively. One *Atoh1^Tom^*;*Cdx::FLPo;Ai65* mouse spinal cord and hindbrain was cleared (male, P30). The hindbrain was imaged as described above (4800 slices). The spinal cord was imaged with a 15x objective with 0.41 μm x 0.41 μm x 2 μm sampling (900 slices). All hindbrain and spinal cord samples were cut to less than 2.2 cm to fit in the imaging chamber. Movies were made in arivis Vision4D 2.12.6. Maximum intensity projections (MIPs) were processed using Fiji (Schindelin et al., 2012).

### Behavioral test - Rotarod

Rotarod was performed at 8-13 weeks of age. All testers were blind to genotype. Mice were acclimated to the testing room for 0.5-1 hr on day of testing. All mice were genotyped for appropriate alleles using previously published protocols for *Atoh1^Cre^*, *Cdx::FLPo*, and *R26^LSL-FSF-TeTx^* (Kim et al., 2009; Yang et al., 2010; Bourane et al., 2015). Mice were placed on an accelerating rotarod (IITC Life Science Series 8, Woodland Hills, CA) from 4 to 40 RPM over 5 min. Four trials were performed each day for two days with at least a 15 min wait time between trials.

### Experimental Design and Statistical Tests

The mossy fiber (MF) to cell body ratio (Fig. 3K) was counted from three cleared spinal cords and hindbrains. Cells bodies in the spinal cord and MFs in the cerebellar cortex were counted from 100 μm MIP images of cleared tissue. The ratio calculated is an estimate given that it is impossible to accurately count all the cell bodies and MF terminals and there are many opportunities for over- or undercounting. For example, although cell bodies and mossy fibers were counted only when they could be discretely identified, some MF terminals might appear as two MF terminals, when they in fact come from the same MF. As an example of undercounting, cell bodies and mossy fibers may overlap in the z axis of the 100 μm MIP and may be counted as one instead of several. Altogether, cell bodies and MFs were counted to get an estimate rather than an exact count of the MF/cell body ratio.

For the mapping of thoracolumbar CC MF terminals in the cerebellum (Fig. 4H-H’’’, J-J’’’, L-L’’’, N, P, R), confocal images of 30 μm cryosections were analyzed in Fiji using the ROI Manager to label individual MF terminals and the SlideSet PlugIn to export the ROIs as a .svg file (Schindelin et al., 2012; Nanes, 2015). These mapped MF terminals were then overlaid on a traced drawing of the confocal image in Adobe Illustrator 2021.

All data and graphs were processed in Microsoft Excel 2015 and GraphPad Prism 9. Details of the number of sections counted and sex of the mice are given in the Results section. Mean ± SEM is reported throughout the manuscript. Statistical tests used are detailed in the Results and/or Figure Legends.

## Results

### Clarke’s column is the major direct spinocerebellar pathway in mice

To assess the spinocerebellar system in mice, we identified genetic tools that reproducibly label spinal cord neurons and evaluated their contribution to the spinocerebellar system using a combination of retrograde and anterograde tracing. Previously, we found that the *Atoh1*-expressing progenitor population that makes dorsal interneuron 1 (dI1) neurons, although implicated in making spinocerebellar neurons developmentally (Bermingham et al., 2001; Gowan et al., 2001; Sakai et al., 2012), rarely made CC neurons, which are the major source of the DSCT (Fig. 1A, note the absence of TOM^+^ cells in the CC area of *Atoh1*-lineage traced neurons, *Atoh1^Cre^;R26^LSL-tdTom^* (Ai14), abbreviated *Atoh1^Tom^*)(Madisen et al., 2010; Yang et al., 2010; Yuengert et al., 2015). Therefore, we sought to identify the progenitor population that gives rise to CC neurons. Evidence from spinal cord development suggested that the neighboring *Neurog1*-expressing progenitor population that makes dI2 neurons also project to the cerebellum (Avraham et al., 2009; Sakai et al., 2012). Because there are no uniquely specific molecular markers for the dI2 population, we traced the lineage of the entire *Neurog1* population, which includes dI2 neurons and ventral domains, using a transgenic *Neurog1BAC-Cre* strain crossed to a *R26^LSL-LacZ^* reporter mouse (Fig. 1A)(Soriano, 1999; Quinones et al., 2010). Large CC cells residing in the medial thoracic spinal cord colocalize with vesicular glutamate transporter 1 (*Vglut1*) mRNA, a marker for CC (Fig. 1A, B)(Llewellyn- Smith et al., 2007; Malet et al., 2013; Yuengert et al., 2015). Therefore, CC neurons come from a *Neurog1*-lineage (dI2 or ventral lineages), but not the *Atoh1*-lineage.

**Figure 1.**
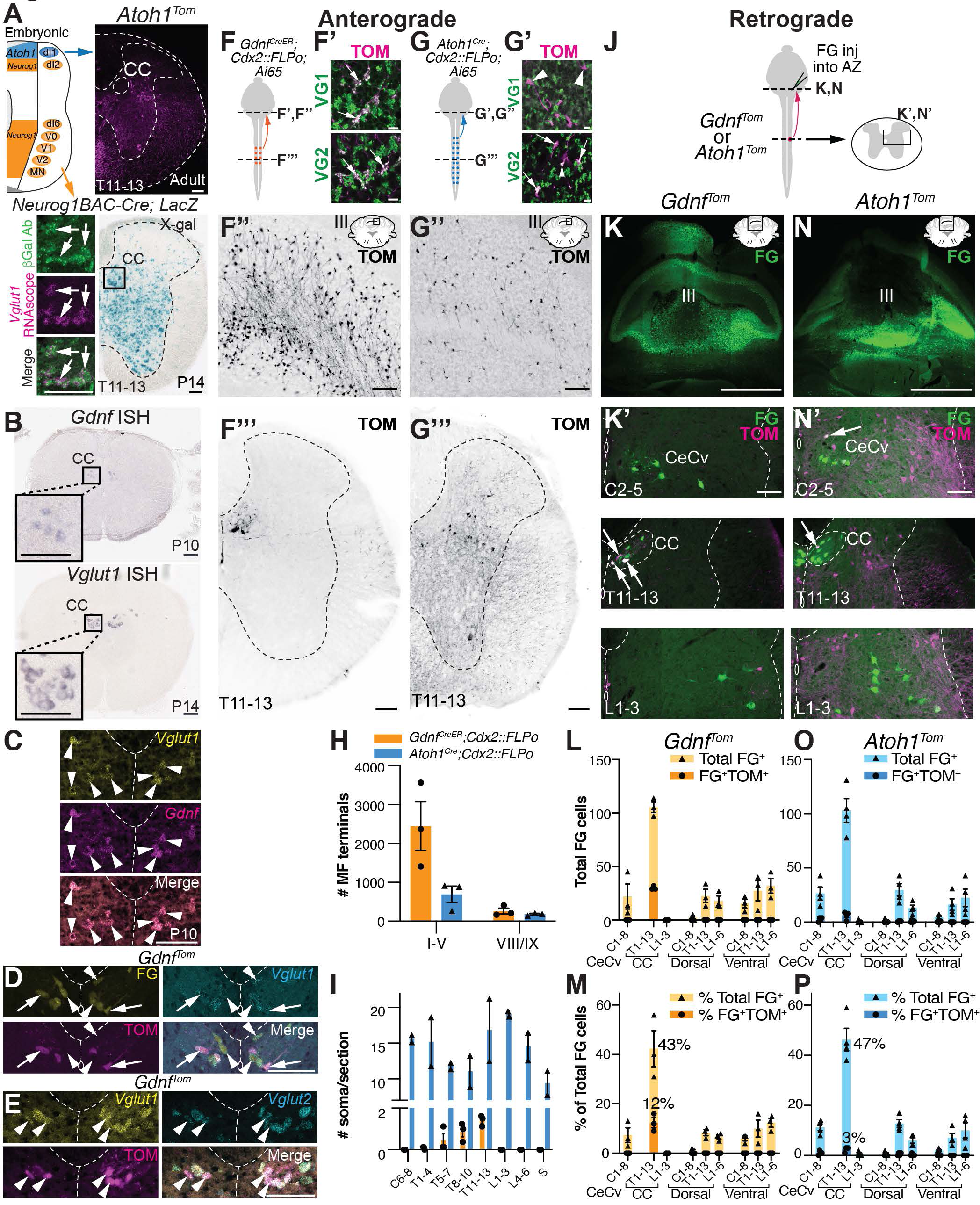
Clarke’s column (CC) is the major direct spinocerebellar pathway in mice. (A) Lineage tracing of *Neurog1*-expressing progenitors (*Neurog1BAC-Cre* crossed to *R26^LSL-LacZ^*) in the neural tube identifies large CC neurons in the thoracic spinal cord (box in X-gal stain). β-Gal expressing cells (green) colocalize with the CC marker, *Vglut1* mRNA (magenta, arrows). *Atoh1*-lineage neurons (*Atoh1^Tom^*) reside lateral and ventral to CC. (B) CC is marked by expression of *Gdnf* and *Vglut1* mRNA. (C) *Gdnf* and *Vglut1* mRNA colocalize in CC neurons at P10 by RNAscope (arrowheads). (D) CC neurons retrogradely labeled with fluorogold (FG) colocalize with *Vglut1* mRNA (FG^+^*Vglut1*^+^, arrowheads). A subset of CC neurons are labeled with *Gdnf^Tom^* (FG^+^TOM^+^*Vglut1*^+^, arrows). (E) CC neurons marked by *Gdnf^Tom^* express both *Vglut1* and *Vglut2* mRNA (arrowheads). (F-I) Comparison of mossy fiber (MF) terminals in the cerebellum of CC and caudal *Atoh1*-lineage neurons. Diagram of CC and caudal *Atoh1*- lineage neurons labeled with tdTomato (TOM)(F, G). MF terminals in the cerebellum of CC neurons are VGLUT1^+^ (VG1) and VGLUT2^+^ (VG2) (F’, arrows). MF terminals of caudal *Atoh1*-lineage neurons are VG2^+^ (G’, arrows), but VG1^-^ (G’, arrowheads). Fewer MF terminals are seen in the cerebellum from caudal *Atoh1*-lineage neurons (G’’) than CC neurons (F’’). Representative thoracic sections of CC and caudal *Atoh1*-lineage neurons (F’’’, G’’’). Quantitation of MF terminals in the vermis of lobules I-V and VIII/IX (H) and of the number of soma in the spinal cord (I) for CC and caudal *Atoh1*-lineage neurons. (J- P) Comparison of spinocerebellar neurons retrogradely labeled with FG in mice with CC (*Gdnf^Tom^*) or *Atoh1*-lineage (*Atoh1^Tom^*) neurons labeled with tdTomato (TOM). Diagram of FG cerebellar injections into the anterior zone (AZ, lobules I-V) of either *Gdnf^Tom^* or *Atoh1^Tom^* mice to retrogradely label direct spinocerebellar projections (J). FG injection in the cerebellum (K, N, green) retrogradely labels central cervical (CeCv) cells in the cervical spinal cord (K’, N’, upper panels), CC in the thoracic spinal cord (K’, N’, middle panels), and neurons in other areas of the spinal cord (K’, N’, lower panels). Retrogradely labeled CC neurons (green) colocalizes with the genetic label for CC (*Gdnf^Tom^*)(K’, middle panel, FG^+^TOM^+^, arrows), but only occasionally colocalizes with *Atoh1*-lineage (*Atoh1^Tom^*) neurons in CeCv or CC (N’, arrows). Quantitation of the total number of FG^+^ cells (L, O) and percentage of total FG^+^ cells (M, P) in a given region of the spinal cord (light orange or light blue) with the total number or percentage of FG^+^TOM^+^ cells superimposed (dark orange or dark blue) for CC and caudal *Atoh1*-lineage neurons is shown. Spinal cords were divided into cervical (C), thoracic (T), and lumbar (L) areas. The CeCv and CC areas are delineated separately with all other cells categorized based on their C, T, or L location and whether they were dorsal or ventral to the central canal. Spinocerebellar cells dorsal or ventral to the central canal are generally not labeled by *Gdnf^Tom^* or *Atoh1^Tom^* (L, M, O, P). Abbrev: P, postnatal. Scale bars: 1 mm (K, N), 100 μm (A-E, F’’-F’’’, G’’-G’’’, K’, N’), 10 μm (F’,G’). Results in graphs presented as mean ± SEM.

Next, we further characterized molecular tools that label CC neurons. As previously described, CC neurons are marked by glial derived nerve growth factor (*Gdnf*) and *Vglut1* mRNA (Fig. 1B)(Llewellyn- Smith et al., 2007; Hantman and Jessell, 2010; Malet et al., 2013). We found that *Gdnf* is transiently expressed from E18.5-P10 (data not shown) and that *Gdnf* and *Vglut1* mRNA completely overlap at P10 (Fig. 1C, arrowheads). Furthermore, CC neurons that are retrogradely labeled with Fluorogold (FG) injected into the anterior zone (AZ, lobules I-V) of the cerebellum, colocalized with *Vglut1* mRNA (Fig. 1D, arrowheads). A subset of CC neurons is labeled using a *Gdnf^IRES2-CreERT2/+^* mouse line crossed to a CRE-dependent tdTomato reporter (Ai14) (abbreviated *Gdnf^Tom^* from here on)(Fig. 1D, arrows)(Hantman and Jessell, 2010; Cebrian et al., 2014). Lastly, we found that CC neurons also express vesicular glutamate transporter 2 (*Vglut2*)(Fig. 1E, arrowheads). We used the *Gdnf^IRES2-CreERT2/+^* mouse line (abbreviated *Gdnf^CreER^* from here on) for the remainder of the study due to its specific labeling of CC neurons.

To understand the relationship of the direct spinocerebellar system to our two CRE mouse lines (*Gdnf^CreER^* and *Atoh1^Cre^*), we used both anterograde (Fig. 1F-I) and retrograde (Fig. 1J-P) tracing strategies to and from the cerebellum. To identify ascending projections from the spinal cord, we used an intersectional strategy to restrict labeling of neurons to regions caudal of cervical 4 (C4)(*Cdx2::FLPo*)(Bourane et al., 2015). As expected, we found that CC neurons (*Gdnf^CreER^; Cdx2::FLPo; Ai65*) terminate as mossy fibers (MF) in the AZ of the cerebellum and express VGLUT1^+^ (VG1) and VGLUT2^+^ (VG2) presynaptic markers (Fig. 1F-F’’’). However, labeling of caudal *Atoh1*-lineage neurons (*Atoh1^Cre^*; *Cdx2::FLPo*; *Ai65*) had very few MF terminals in the AZ, although they were VG1^-^ and VG2^+^, consistent with a non-Clarke’s spinocerebellar population and a previous report that spinocerebellar neurons are mainly VG2^+^ with some VG1^+^ MF terminals (Gebre et al., 2012; Yuengert et al., 2015)(Fig. 1G-G’’’; Quantitated in Fig. 1H, in the AZ, 2447 ± 624 *Gdnf^CreER^; Cdx2::FLPo; Ai65* (n=3, 2 females (F):1 male (M)) vs. 690 ± 213 *Atoh1^Cre^*; *Cdx2::FLPo*; *Ai65* (n=3, 1F:2M), t-test p=0.056, 3 comparable sections/n; MF terminals in lobules VIII/IXa are similar, 262 ± 75 *Gdnf^CreER^; Cdx2::FLPo; Ai65* vs. 182 ± 21 *Atoh1^Cre^*; *Cdx2::FLPo*; *Ai65,* 1 comparable section/n; ages P27-67). The difference in MF terminals is striking given that *Atoh1*-lineage neurons have many more soma per section throughout the rostral- caudal axis compared to CC neurons (Fig. 1I, *Gdnf^CreER^; Cdx2::FLPo; Ai65* (n=3, 2F:1M, 4 sections counted per region per n), *Atoh1^Cre^*; *Cdx2::FLPo*; *Ai65* (n=2, 1F:1M, 3-4 sections counted per region per n)), suggesting there are many more MF terminals per CC soma in the spinal cord compared to caudal *Atoh1*-lineage neurons.

Consistent with our anterograde tracing findings, we found that retrograde labeling of spinocerebellar neurons colocalized with CC neurons, but few *Atoh1*-lineage neurons (Fig. 1J-P). FG was injected into the cerebella of *Gdnf^Tom^* or *Atoh1^Tom^* mouse strains targeting the AZ where spinocerebellar neurons are known to project (Fig. 1 J, K, N)(Arsenio Nunes and Sotelo, 1985; Bosco and Poppele, 2001; Reeber et al., 2011). Overall, FG retrograde tracing from all injections were similar with 261 ± 39 total FG^+^ cells in injections of *Gdnf^Tom^* (n=3, 1F:2M, ages P38-40, counts from 5 sections per spinal cord region per n) and 228 ± 32 total FG^+^ cells in injections of *Atoh1^Tom^* (n=4, 3F:1M, ages P39-40, counts from 5 sections per spinal cord region per n)(Fig. 1L, O shows total FG cells in each spinal cord region). Overall, CC is the most abundantly labeled spinocerebellar projection making up 43- 47% of all retrogradely-labeled FG neurons in the spinal cord (Fig. 1M, P). The next most abundant areas of spinocerebellar neurons along the rostral-caudal axis are those in the central cervical (CeCv) nucleus (Cummings and Petras, 1977; Wiksten, 1987; Popova et al., 1995) and cells dorsal or ventral of the central canal in the thoracolumbar areas (Fig. 1K’, N’, top and lower panels; L-M, O-P), which correspond to LV-SCT, LVII-SCT, LVIII-SCT, and spinal border cells (Baek et al., 2019). Notably, spinocerebellar neurons in the CeCv and thoracolumbar areas (excluding CC) are rarely colabeled with *Gdnf^Tom^* or *Atoh1^Tom^* neurons (Fig. 1L-M, O-P) indicating that genetic labels for these spinocerebellar neurons and their developmental origins have yet to be determined. *Gdnf^Tom^* makes up 12 ± 2% of the FG^+^ CC neurons out of all FG^+^ neurons in the spinal cord and therefore labels ∼ 29 ± 1% of CC neurons. The remaining approximately 70% of unlabeled CC neurons in the *Gdnf^Tom^* line could be due to the restricted time point at which tamoxifen was injected (P7-P8), incomplete CRE recombination, or represent a unique subset of CC neurons. *Atoh1^Tom^* makes up 3 ± 1% of the FG^+^ CC neurons out of all FG^+^ neurons in the spinal cord and therefore labels ∼ 6 ± 1% of CC neurons, consistent with our previous findings that *Atoh1*-lineage neurons make up very few CC neurons (Yuengert et al., 2015). Significantly, only 4 ± 1% of all spinocerebellar (FG^+^) neurons projecting to the AZ in the entire spinal cord are *Atoh1*- lineage neurons (FG^+^TOM^+^). Altogether, our data suggest that CC makes up a majority of spinocerebellar neurons projecting directly to the AZ, while *Atoh1*-lineage neurons make up very few direct spinocerebellar neurons. Using our genetic tools, we then proceeded to determine the precise anatomy of both CC and *Atoh1*-lineage projections.

### Anatomical trajectories of CC neurons

We used the *Gdnf^Tom^* line to meticulously trace axonal trajectories of CC neurons to the cerebellum and found that the axons cross within the cerebellum and terminate almost exclusively as MFs on granule cells. We found that CC MF terminals in the cerebellum terminate in the vermis of lobules II-V, VIII, IXa, and the copula pyramidis (Cop) (Fig. 2A-C), consistent with the termination locations of spinocerebellar neurons from previous pan-anterograde tracing studies (Arsenio Nunes and Sotelo, 1985; Bosco and Poppele, 2001; Apps and Hawkes, 2009; Reeber et al., 2011). In addition, the three parasagittal stripes in lobule III on both sides of the midline closely matched those found in anterograde tracing studies from the thoracic and lumbar spinal cord (Fig. 2A, B)(Ji and Hawkes, 1994; Reeber et al., 2011). However, while CC axons are known to travel rostrally ipsilaterally in the lateral funiculus of the spinal cord (Oscarsson, 1965 and see Movie 1), we found that several axons appeared to cross the midline within the cerebellum (Fig. 2D, E), suggesting that CC axons terminate both ipsilaterally and contralaterally in the cerebellar cortex, which has been seen in single-cell reconstructions (Luo et al., 2018). To test whether CC axons from a given CC cell terminates on both sides of the cerebellum, we injected two different Cholera Toxin subunit B (CTB)-conjugated fluorophores (CTB-488 and CTB-647) into the left and right sides of the cerebellum (Fig. 2F, G). We found retrogradely labeled cells in CC of the spinal cord that took up both tracers (Fig. 2H, arrows and arrowheads), some of which were colabeled with *Gdnf^Tom^* (Fig. 2H, arrows). Approximately 30% of the terminals from CC neurons that innervate the injection area are from the contralateral side (% ipsi and contra CTB-488/total CTB-488: 74 ± 4% and 26 ± 4%, respectively; % ipsi and contra CTB-647/total CTB-647: 70 ± 5% and 30 ± 5%, respectively, n=3, 2F:1M, 8 sections/n, age P65-95). This could mean that only 30% of CC neurons cross or that all of them cross, but only some CC neurons have enough terminations on the contralateral side to allow for sufficient CTB uptake. Therefore, CC neurons project ipsilaterally within the spinal cord, but send collaterals to both ipsilateral and contralateral sides within the cerebellum.

**Figure 2.**
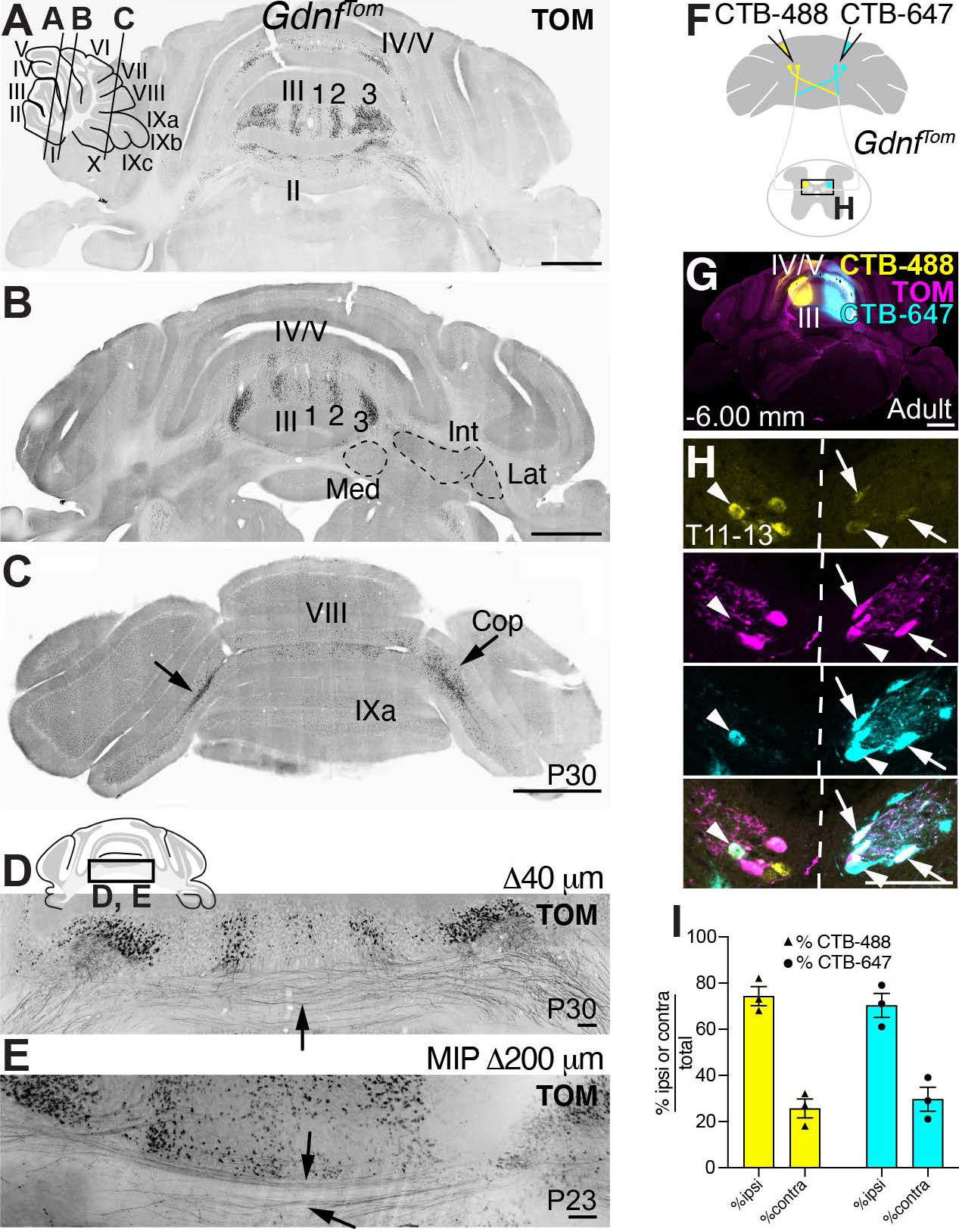
CC mossy fibers terminate ipsilaterally and contralaterally in the cerebellar vermis. **(A-**C) Coronal sections from *Gdnf^Tom^* mice reveal CC mossy fiber (MF) terminals (TOM^+^) in lobules II-V, VIII, IXa, and the copula (C, Cop, arrows). Parasagittal stripes (1, 2, 3) in lobule III are apparent. (D-E) Some CC axons (TOM^+^) cross the midline (D, arrow, cryosection, and E, arrows, cleared sample, 200 μm maximum intensity projection (MIP)). (F) Diagram of dual CTB-488 and CTB-647 injections in *Gdnf^Tom^* mice. (G) Coronal section showing the injection site of CTB-488 and CTB-647. (H) CC neurons are co- labeled with the fluorescent CTB injected on the ipsilateral side as well as the fluorescent CTB injected on the contralateral side (CTB-488^+^CTB-647^+^, arrows and arrowheads,). Some cells also colocalize with the *Gdnf^Tom^* genetic label for CC (TOM^+^CTB-488^+^CTB-647^+^, arrows) and some do not (TOM^-^CTB- 488^+^CTB-647^+^, arrowheads). (I) Quantitation of the percentage of ipsilaterally or contralaterally labeled CTB^+^ cells out of all CTB^+^ cells labeled in the spinal cord with a particular CTB fluorophore (mean ± SEM). Abbrev: Med, Medial; Int, Interpositus; Lat, Lateral. Scale bars: 1 mm (A, B, C, G), 100 μm (D, E, H).

Strikingly, we found that CC axons do not make significant axon collaterals within the spinal cord, medulla, or to the cerebellar nuclei, a feature typical of other MF tracts, but has been ambiguous for the spinocerebellar system (Fig. 3A-H’)(Matsushita and Ikeda, 1970; Matsushita and Gao, 1997; Mogensen et al., 2017). In three separate samples, we found no axon terminations in the Medial (Med), Interpositus (Int), or Lateral (Lat) cerebellar nuclei (CN). Areas near the cerebellar nuclei that had TOM^+^ signal came from axons of passage and not synaptic terminations (TOM^+^ axons are VG1^-^ or VG2^-^ (Fig. 3A’, A’’, D’, D’’)). Furthermore, we find no seeming axon collaterals within the spinal cord (Movie 1), nor do we find significant synaptic terminations in the LRt or nucleus X, as was reported previously for some CC neurons (Fig. 3G-H’)(Luo et al., 2018). Some synaptic terminals can be seen in the LRt (Fig. 3G’, arrow), but the axons mainly appear to be coursing by the LRt. In summary, our genetic studies of CC neurons show that these glutamatergic neurons terminate bilaterally in the cerebellar vermis, but do not make significant axon collaterals to the spinal cord, medulla, or cerebellar nuclei.

**Figure 3.**
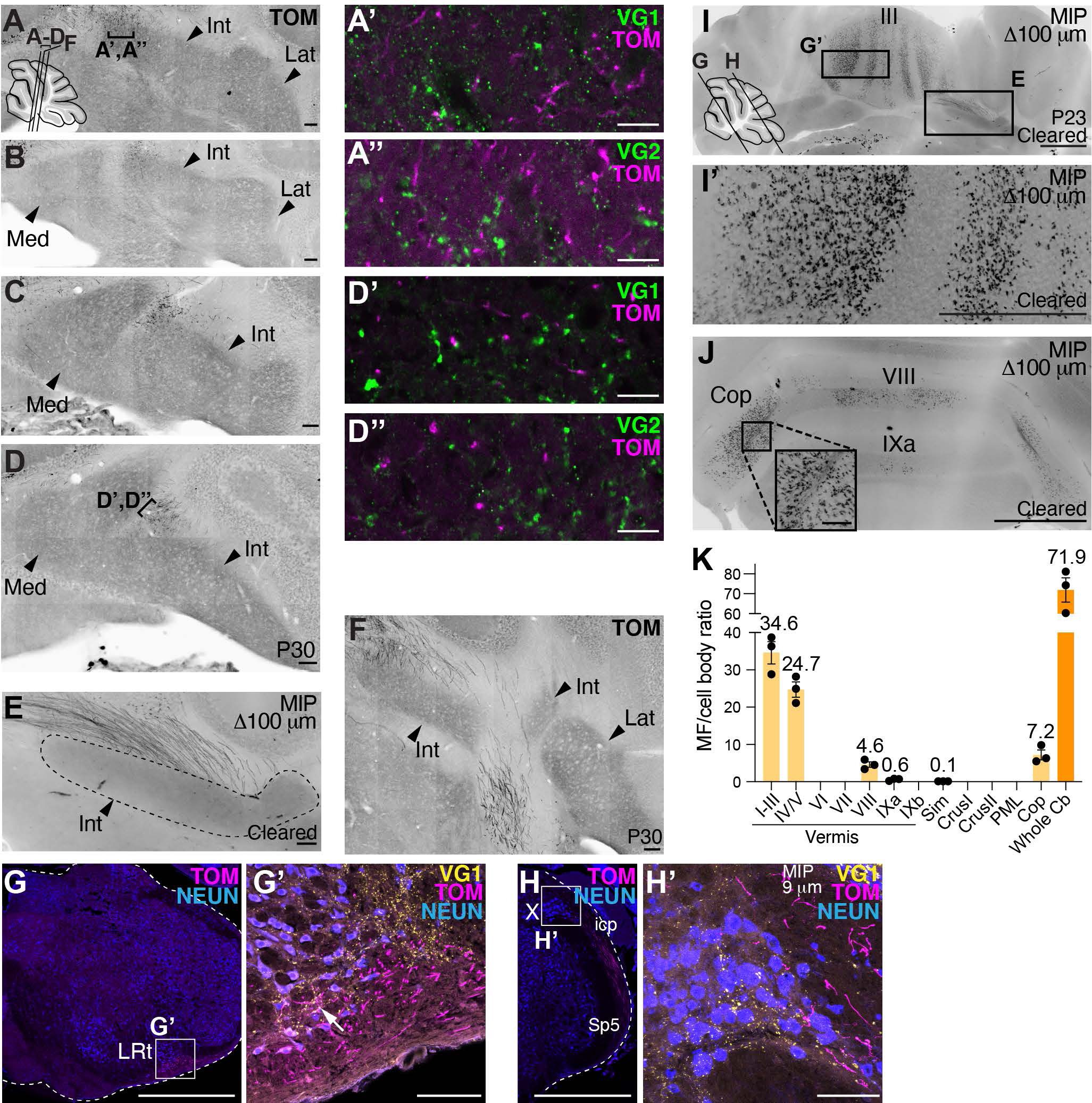
CC neurons do not collateralize to the medulla or cerebellar nuclei (CN), but arborize extensively in the cerebellar cortex. (A-D’’) Almost no CC *Gdnf^Tom^* axons enter or are near the CN (arrowheads, Med, Int, Lat)(A, B, C, and D are successive sections 160 μm apart). Areas of TOM^+^ signal near CN do not colocalize with presynaptic markers VG1 (A’, D’) or VG2 (A’’, D’’) indicating these are axons of passage and not presynaptic terminals. (E) CC axons (TOM^+^) avoid the CN in another *Gdnf^Tom^* mouse whose cerebellum was cleared (100 μm MIP). (F) CC axons (TOM^+^) avoid the CN (40 μm cryosection) in another example *Gdnf^Tom^* mouse. Images are from two female (A-E) and one male (F) mice (n=3). (G-H’) Only a few axonal terminations are seen in the LRt (G-G’, arrow) and none in nucleus X (H-H’)(verified in n=3 mice, representative sections shown). (I-K) Example images of 100 μm MIP from a *Gdnf^Tom^* cleared cerebellum. CC MF terminals (TOM^+^) are seen in II-V, VIII, IXa, and Cop. (K) Quantification of the MF/cell body ratio (n=3 mice, one *Gdnf^Tom^* and two *Gdnf^CreER^; Cdx2::FLPo; Ai65* mice). Overall the whole cerebellum (Cb) has an estimated 71.9 MF terminals per CC cell body in the spinal cord (orange bar, mean ± SEM). Most MF terminals from CC cells terminate in I-III, IV/V, VIII/IXa, and Cop (light orange bars). Abbrev: Med, Medial; Int, Interpositus; Lat, Lateral. Lateral reticular nucleus, LRt; X, Nucleus X; icp, inferior cerebellar peduncle; Sp5, spinal trigeminal tract. Scale bars: 1 mm (G, H, I, I’, J), 100 μm (A-F, G’, H’, J inset), 10 μm (A’-A’’, D’-D’’).

### Diversification of proprioceptive information through CC neurons

To obtain a three-dimensional view of CC trajectories, we chemically cleared the spinal cords and hindbrains of *Gdnf^Tom^* and *Gdnf^CreER^;Cdx2::FLPo*; *Ai65* mice (Movies 1-2). Because *Gdnf* is also expressed in smooth and skeletal muscle (Trupp et al., 1995; Suzuki et al., 1998; Rodrigues et al., 2011), they are prominently labeled with TOM in these samples. In the spinal cord (Movie 1), the CC soma can be seen straddling the midline, while their axons extend to the lateral funiculus (LF) where they make a 90° turn heading rostrally to the cerebellum. Axons in the inferior cerebellar peduncle are seen traveling directly to the cerebellum (Movie 2).

A feature that was readily apparent from the cleared specimens was the sheer number of MF terminals in the cerebellum indicating an immense diversification of proprioceptive information coming from CC axons (Fig. 3I-K). We counted the number of MF terminals per CC soma in the cerebella and spinal cords of the three cleared samples. From these counts, we estimate that there are 71.9 ± 6.1 MF terminals in the entire cerebellum for each CC soma (Fig. 3K, orange bar). The MFs terminate largely in vermis I-III (34.6 ± 3.0 MF/soma ratio), IV/V (24.7 ± 2.1), VIII (4.6 ± 0.7), IXa (0.6 ± 0.2), and the copula (7.2 ± 1.3)(n=3, 2F:1M, ages P23-28), consistent with the distribution seen in cryosections (Fig. 2A-C). The large ratio of MFs to CC soma suggests that CC information is widely distributed within the cerebellum. Furthermore, the proprioceptive information coming into CC neurons require surprisingly few CC neurons to relay that information to the cerebellum. We counted a range of 486-816 CC neurons in the three spinal cords, which represents around 29% or less of all CC neurons labeled in the *Gdnf^Tom^* and *Gdnf^CreER^;Cdx2::FLPo*; *Ai65* mouse models for an estimated 1,620 to 2,720 CC neurons in the mouse spinal cord. This suggests that most of the mouse proprioceptive direct spinocerebellar system comes from roughly a couple thousand neurons.

Next, we wanted to test whether CC neurons from a restricted area of the spinal cord terminate in clustered or diverse locations in the cerebellum. If a given CC neuron sends MFs terminals to one discrete localized area of the cerebellum, this would suggest that proprioceptive information exists as a traditional homunculus in the cerebellum. However, if a given CC neuron sends MF terminals to multiple areas of the cerebellar cortex, this would provide an anatomical substrate for the fractured somatotopic map that has been detected electrophysiologically (Shambes et al., 1978), where body parts are represented in discontinuous patches across the cerebellum (Manni and Petrosini, 2004; Apps and Hawkes, 2009). To label CC neurons specifically in the thoracolumbar area, we injected AAV9-Syn-DIO- EGFP into *Gdnf^Tom^* mice (Fig. 4A). Our injections labeled CC neurons on both sides of the spinal cord (Fig. 4B, B’, arrows) and largely the lower thoracolumbar spinal cord (Fig. 4G, I, K, M, O, Q)(counts for GFP^+^ infection in the spinal cord are from 15-16 sections per spinal cord region). We found that thoracolumbar CC neurons targeted multiple lobules (II-V, VIII)(GFP^+^ (green) and GFP^+^TOM^+^ (yellow), Fig. 4H-H’’’, J-J’’’, L-L’’’, N, P, R, coronal and sagittal sections, n=3 each, 1F:2M for coronal sections, 2F:1M for sagittal sections). Although there were discrete areas that did not contain GFP^+^ cells (arrowheads, Fig. 4H-H’’’, J-J’’’, L-L’’’), GFP^+^ cells were found over multiple lobules indicating that CC axonal projections from the thoracolumbar spinal cord terminate throughout the cerebellar cortex, consistent with a discontinuous somatotopic map. Examples of GFP^+^ (green, arrowheads) and GFP^+^TOM^+^ (white, arrows) MF terminals in lobules III and VIII in both coronal and sagittal sections are shown (Fig. 4S, T). In addition to terminating across several lobules, we found multiple examples of single axons terminating at regular intervals (50-80 μm) within a GC layer, which has been reported anecdotally in the literature (Reeber et al., 2011; Houck and Person, 2015; Gilmer and Person, 2017; Luo et al., 2018), indicating that a single CC neuron synapses on several GCs (Fig. 4C-F). Interestingly, an individual GC is approximately 40-50 μm from dendrite to dendrite (Gray, 1961; Eccles et al., 1967; Jakab and Hamori, 1988; Huang et al., 2013). Therefore, individual MF axons from a single CC neuron likely do not synapse on the same GC suggesting that GCs are multimodal encoders at the single cell level (Marr, 1969; Albus, 1971). Overall, we find that CC neurons arborize extensively within the cerebellar cortex, reaching targets over multiple lobules, rather than in clustered locations.

**Figure 4.**
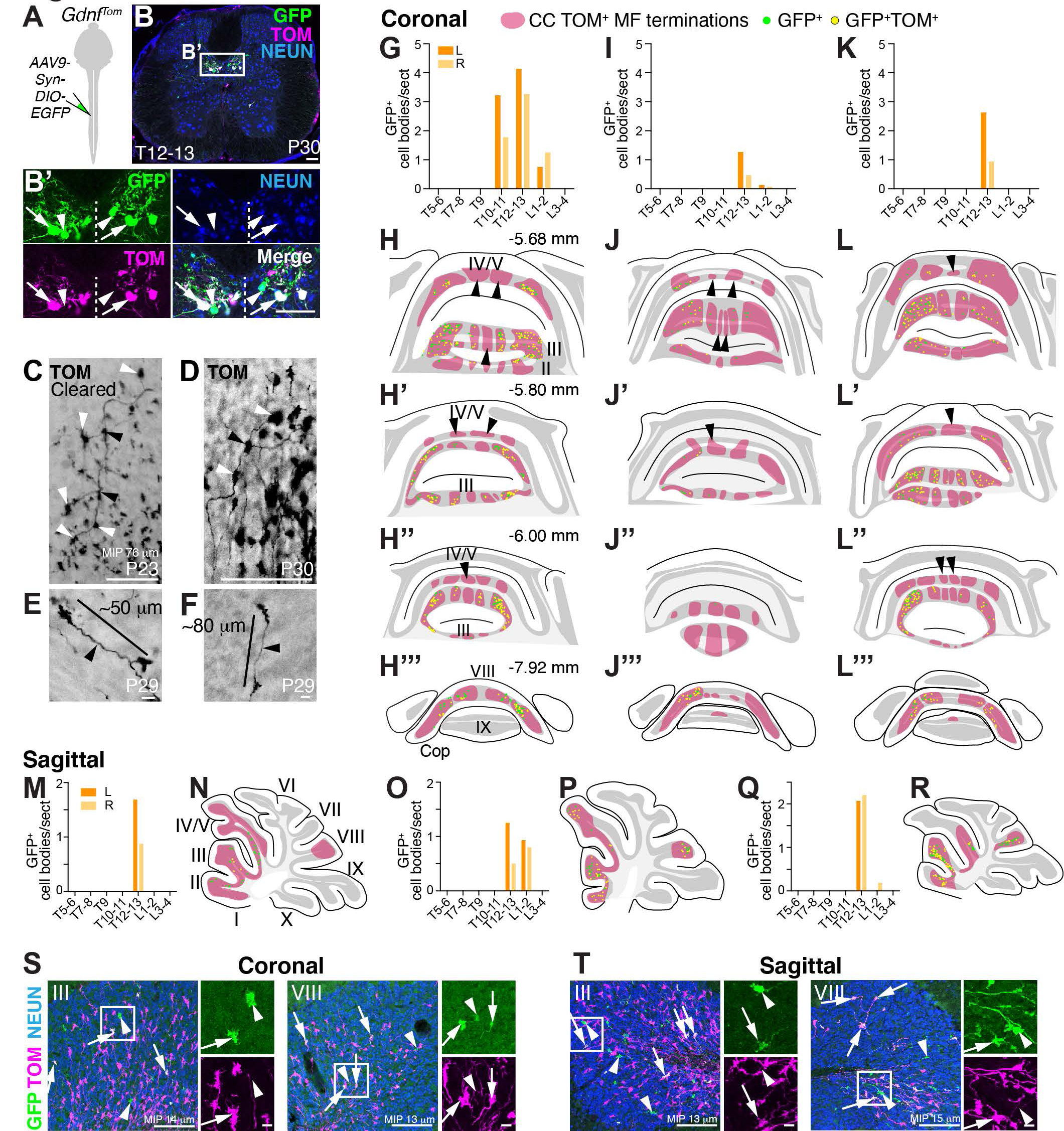
Thoracolumbar CC MFs send diverse projections to multiple lobules. (A-B’) Spinal cord injections of AAV9-Syn-DIO-EGFP at lower thoracic levels into *Gdnf^Tom^* mice labels CC neurons on both sides of the spinal cord (B’, GFP^+^TOM^+^ (arrows), GFP^+^ only (arrowheads)). (C-F) Examples of individual MF axons and terminals from three mice: a cleared female mouse sample (C, 76 μm maximum intensity projection (MIP)), one female mouse (D, 40 μm cryosection), and one male mouse (E, F, 40 μm cryosection). Axons appear to have branching points (black arrowheads) and regularly spaced MF terminals (white arrowheads)(C-D). MF terminals from an individual axon are spaced 50-80 μm apart (E- F). (G-L’’’) Thoracolumbar injections whose MF terminations were analyzed in coronal sections. Distribution of GFP^+^ cells in the spinal cord on left (orange) and right (light orange) sides (G, I, K, M, O, Q). Schematics of coronal cerebellar sections of the spinal cord injections (G, I, K) indicating the location of CC MF terminations (H-H’’’, J-J’’’, L-L’’’, TOM^+^, red areas, respectively). Schematics of sagittal cerebellar sections of the spinal cord injections (M, O, Q) indicating the location of CC MF terminations (N, P, R, red areas, respectively). The subset of CC MF terminations that are from the lower thoracic- lumbar region (GFP^+^, green, and GFP^+^TOM^+^, yellow) are spread over multiple lobules (II-V, VIII). Certain CC MF termination regions do not have thoracolumbar CC neuronal projections (red areas, arrowheads with an absence of any GFP^+^ terminations). (S, T) Examples of CC MF terminations (GFP^+^TOM^+^, arrows, and GFP^+^-only arrowheads) in lobules III and VIII from 13-15 μm maximum intensity projections (MIP). Scale bars: 100 μm (B, B’, C, D, S, T); 10 μm (E, F).

### Atoh1-lineage neurons make the spino-LRt and spino-olivary tracts

Given that *Atoh1*-lineage neurons made few direct spinocerebellar neurons, we sought to identify where in the hindbrain *Atoh1*-lineage axons project. We pursued an intersectional genetic strategy to restrict somatic labeling to caudal *Atoh1*-lineage neurons (*Atoh1^Cre^; Cdx2::FLPo; Ai65*)(Fig. 5A). Most prominently, we found dense projections of caudal *Atoh1*-lineage neurons in the lateral reticular nucleus (LRt) and inferior olive (IO) (Fig. 5E-H’’). To identify whether caudal *Atoh1*-lineage axons synapse on localized areas of the LRt and IO, we injected FG into the cerebellar AZ (Fig. 5A-B), which retrogradely labeled LRt MF and IO CF cell bodies. We found that caudal *Atoh1*-lineage axons target almost the entirety of the LRt and restricted areas of the IO (dorsal fold of the dorsal accessory olive (dfDAO), dorsal accessory olive (DAO), subnucleus a of the caudal medial accessory olive (cMAO^a^), subnucleus b of cMAO (cMAO^b^)(Fig. 5E’-H’). Consistent with our findings, anterograde tracing in rats reports that the spino-olivary tract terminates in the DAO, cMAO^a^, and cMAO^b^ (Swenson and Castro, 1983a; Matsushita et al., 1992; Oldenbeuving et al., 1999). Caudal *Atoh1*-lineage axons colocalize with the presynaptic marker VG2 and are in close apposition to FG labeled neurons in the LRt and IO indicative of synaptic connections (arrows, Fig. 5I-J, Movies 5 and 6). Axon terminations in the LRt and IO were verified in four caudal *Atoh1*-lineage mice. Moreover, TOM^+^ terminals in a cleared brain of *Atoh1^Cre^; Cdx2::FLPo; Ai65* mice are quite dense in the LRt and IO (Movie 4). Altogether, we find that spinal cord *Atoh1*-lineage neurons make the spino-LRt and spino-olivary tracts.

**Figure 5.**
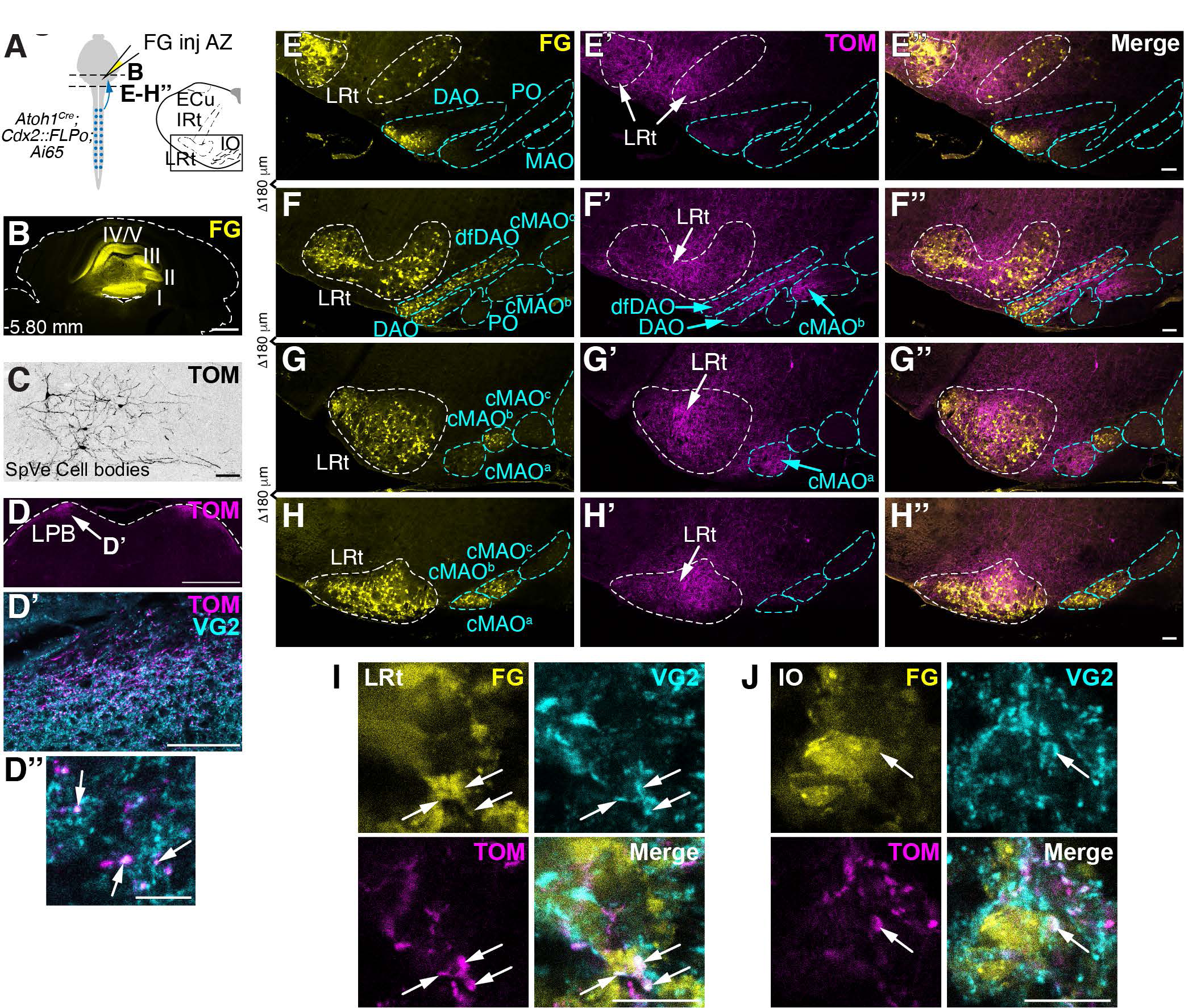
Spinal cord *Atoh1*-lineage neurons make spino-LRt and spino-olivary pathways. (A) Schematic of FG injections into the AZ of mice to identify LRt and IO neurons in the medulla. Axons of caudal *Atoh1*-lineage neurons are genetically labeled (*Atoh1^Cre^; Cdx2::FLPo; Ai65* mice). (B) FG injected into lobules I-V. (C) Sparse cell bodies in the SpVe are detected in *Atoh1^Cre^; Cdx2::FLPo; Ai65* mice. (D- D’’) TOM^+^ terminals seen in the lateral parabrachial (LPB) nucleus (D) are VG2^+^ (D’’, arrows). (E-H’’) A high density of caudal *Atoh1*-lineage axons from the spinal cord are found in the LRt as well as areas of the IO (dfDAO, DAO, cMAO^a^, cMAO^b^). (I-J) Caudal *Atoh1*-lineage axon terminals (TOM^+^, magenta) expressing the presynaptic VG2 marker (cyan) are closely apposed to retrogradely labeled FG^+^ cells in the LRt (I) and IO (J)(arrows). Axonal terminations in the LRt and IO were verified in n=4 mice. Representative sections shown in E-H’’. Abbrev: ECu, external cuneate nucleus; IO, inferior olive; cMAO^a^, subnucleus a of caudal medial accessory olive; cMAO^b^, subnucleus b of caudal medial accessory olive; cMAO^c^, subnucleus c of caudal medial accessory olive; MAO, medial accessory olive; DAO, dorsal accessory olive; dfDAO, dorsal fold of the DAO; PO, principal olive; IRt, intermediate reticular nucleus; LRt, lateral reticular nucleus; SpVe, spinal vestibular nucleus. Scale bars: 1 mm (B, D), 100 μm (C, D’, E-H’’), 10 μm (D’’, I, J).

In the caudal *Atoh1*-lineage mice, a few places of expression are worth noting. First, the sensory neurons misexpress tdTomato and their axons can be seen terminating in the dorsal horn of the cleared spinal cord (Movie 3). *Atoh1* is not known to be expressed in sensory neurons and we do not see sensory neurons labeled in *Atoh1^P2A-FLPo^* mice, whose expression overlaps quite well with the *Atoh1^Cre^* knockin mouse for spinal cord interneurons (data not shown and Ogujiofor et al., 2021). Therefore, the sensory neuron labeling is due to misexpression of the CRE recombinase in sensory neurons. As a result, axons from sensory neurons going to the gracile, cuneate, and external cuneate nuclei can be seen in the cleared brain of *Atoh1^Cre^; Cdx2::FLPo; Ai65* mice (Movie 4). Second, we find sparse ectopic labeling of *Atoh1*-lineage spinal vestibular (SpVe) soma in the hindbrain (Fig. 5C)(Rose et al., 2009). SpVe neurons send descending projections to the spinal cord (Liang et al., 2015), therefore, we do not expect that this sparse ectopic labeling interferes with our analysis of ascending projections from caudal spinal cord *Atoh1*-lineage neurons. Lastly, we detect some TOM^+^ axons in the lateral parabrachial nucleus (LPB) in *Atoh1^Cre^; Cdx2::FLPo; Ai65* mice (TOM^+^VG2^+^, Fig. 5D-D’). Further studies are needed to determine exactly from where in the spinal cord these LPB projections originate and if they are truly *Atoh1*-lineage neurons or ectopic expression.

### Spino-LRt and spino-olivary axonal projections originate from cervical Atoh1-lineage neurons

We pursued both retrograde and anterograde tracing strategies to determine which caudal *Atoh1*- lineage neurons contribute to the spino-LRt and spino-olivary tracts. In four different FG injections targeting the LRt and IO in *Atoh1^Tom^* mice, we found that the retrogradely labeled *Atoh1*-lineage cells resided mainly in the cervical to upper thoracic areas (Fig. 6A-G, light purple (left side) and light blue (right side) FG^+^TOM^+^ cells quantitated in B, C, E, F; n=4, 3F:1M, 5 sections counted per spinal cord region, 7-9 weeks old). *Atoh1*-lineage neurons have previously been described as clustering into lateral and medial populations, which are thought to make up ipsilaterally- and contralaterally-projecting populations, respectively (Bermingham et al., 2001; Wilson et al., 2008; Yuengert et al., 2015). Retrogradely labeled neurons from the LRt and IO colocalized with both the lateral (Fig. 6D’, G’) and medial (Fig. 6G’’) *Atoh1*-lineage neurons. Because of the spread of the FG to both the LRt and IO, the precise ipsilateral or contralateral projections from either tract was not clear, but *Atoh1*-lineage neurons both ipsilateral (Fig. 6G’) and contralateral (Fig. 6G’’) to the injection site were retrogradely labeled. Given the large number of neurons retrogradely labeled from the LRt and IO that are not *Atoh1*-lineage suggests that there are many other progenitor domains and cell types that contribute to these tracts (Alstermark and Ekerot, 2013; Azim et al., 2014; Pivetta et al., 2014; Jiang et al., 2015; Choi et al., 2020).

**Figure 6.**
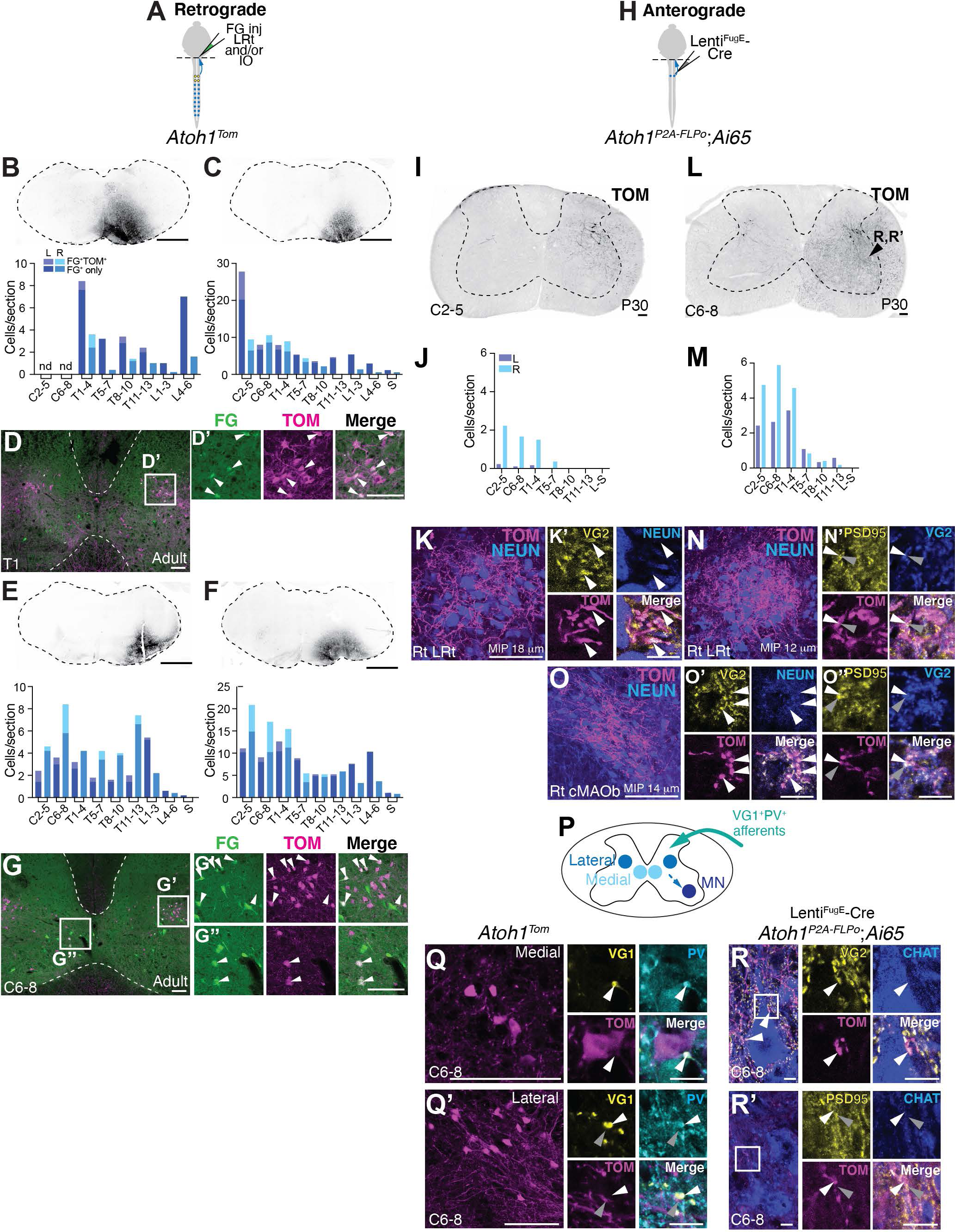
Cervical spinal cord *Atoh1*-lineage neurons project to the LRt and IO. (A) Schematic of FG injections into the right LRt and IO of *Atoh1^Tom^* mice. (B-G) Retrogradely labeled *Atoh1*-lineage neurons reside mainly in the cervical to upper thoracic levels. Four injections targeting the LRt and IO in the medulla are shown (B, C, E, F – upper panels). The spinal cords of these four injections have dual labeled cells (FG^+^TOM^+^) mainly in the cervical to upper thoracic regions (light purple - left side, light blue right side) with many other FG^+^ only cells elsewhere in the spinal cord (dark purple – left side, dark blue right side). Representative spinal cord sections of the injection in C is shown in D-D’ and the injection in F is shown in G-G’’. *Atoh1*-lineage neurons in both the medial (G’’) and lateral (D’ and G’) clusters are retrogradely labeled with FG (arrowheads). (H) Schematic of dual recombinase anterograde tracing. Lenti^FugE^-Cre was injected into the right cervical spinal cord of *Atoh1^P2A-FLPo^*; *Ai65* mice. (I-O’’) Anterograde tracing of *Atoh1*-lineage neurons finds axons in the LRt and IO. Injection of the right spinal cord (I) where mostly neurons on the right side are labeled (J) has axonal projections to the right LRt, which express the presynaptic marker VG2 (K-K’, arrowheads). Injection of the right spinal cord (L) where *Atoh1*-lineage neurons on both the left and right are labeled (M) has axonal projections to both the right LRt and IO, specifically the cMAOb (N, O). The *Atoh1*-lineage axonal terminals (TOM^+^) express the presynaptic marker VG2 (white arrowheads) and are closely apposed to postsynaptic PSD95^+^ punta (grey arrowheads)(N’, O’, O’’). (P) Schematic of inputs and local outputs of cervical *Atoh1*-lineage neurons. (Q- Q’) Synaptic terminals of proprioceptive afferents (VG1^+^PV^+^, white arrowheads) are closely apposed to TOM^+^ signal near medial and lateral *Atoh1*-lineage neurons. (R-R’) Axons of cervical *Atoh1*-lineage neurons synapse on motor neurons (VG2^+^TOM^+^ puncta closely apposed to PSD95^+^ puncta on CHAT^+^ motor neurons, white and grey arrowheads). CHAT^+^ neurons in R-R’ were imaged from the area of the spinal cord indicated in L. Abbrev: nd, not determined. Scale bars: 100 μm (B, C, D-D’, E, F, G-G’’, I, K, L, O, Q, Q’), 10 μm (K’, N’, O’-O’’, Q-Q’ high magnification, R-R’ high magnification).

To identify the axonal projection targets of cervical *Atoh1*-lineage neurons, we pursued an intersectional strategy injecting Lenti^FugE-Cre^ into the cervical area of *Atoh1^P2A-FLPo^* mice crossed to an intersectional tdTomato reporter (*Ai65*) (Fig. 6H). In one mouse, the infected neurons were restricted to the right spinal cord (Fig. 6I-J) and in another mouse, the infected cells were on both the left and right sides (Fig. 6 L, M)(n=2 male mice, 8-15 sections counted per spinal cord region, P30). In the mouse with the injection restricted to the right side, prominent axons were seen in the right LRt and the TOM^+^ terminations express VG2 (Fig. 6K-K’), suggesting that minimally, the spino-LRt *Atoh1*-lineage neurons are ipsilaterally-projecting. In the other mouse that had TOM^+^ cells on both sides of the spinal cord, likely due to the virus being taken up by contralaterally-projecting *Atoh1*-lineage neurons, we found axonal terminations in the right LRt and right cMAOb that were VG2^+^ and closely apposed to the postsynaptic density protein (PSD-95^+^), suggesting that these are functional excitatory synapses (Fig. 6N-O’’, TOM^+^VG2^+^, white arrowheads; PSD95^+^, grey arrowheads).

We next examined the local inputs and outputs of cervical *Atoh1*-lineage neurons. We had previously shown that thoracolumbar *Atoh1*-lineage neurons receive proprioceptive input (Yuengert et al., 2015). Here, we found that both medial and lateral *Atoh1*-lineage neurons in the cervical area (TOM^+^) have processes closely apposed to VG1^+^ and Parvalbumin (PV^+^) synapses indicative of proprioceptive afferents (Fig. 6Q-Q’). Similar to our previous results for the thoracolumbar *Atoh1*-lineage neurons (Yuengert et al., 2015), we were able to visualize axo-somatic synapses for the medial population (Fig. 6Q, VG1^+^PV^+^, white arrowhead), but we were unable to identify axo-somatic synapses on the lateral population. We could only find VG1^+^PV^+^ afferents passing near TOM^+^ processes in the vicinity of the lateral soma (Fig. 6Q’, VG1^+^PV^+^, white arrowhead; TOM^+^, grey arrowhead). Lastly, we found that the axons of sparsely labeled cervical *Atoh1*-lineage neurons made synapses on motor neurons (MNs)(Fig. 6R-R’). Altogether, we found that cervical *Atoh1*-lineage neurons can receive proprioceptive input and do indeed project to the LRt and IO as well as locally to MNs in the cervical spinal cord.

### Thoracolumbar Atoh1-lineage neurons project locally

We next examined whether sparsely labeled thoracolumbar *Atoh1*-lineage neurons send axonal projections to the medulla or cerebellum. Using the same intersectional injection strategy except with the Lenti^FugE-Cre^ targeted to the right thoracolumbar spinal cord (light blue), we again found several cell bodies on the contralateral side labeled (Fig. 7B, 786 ± 20 cells, right side vs. 598 ± 223 cells, left side; n=4, 3F:1M, total number of cells infected was estimated from counts of 20% of each spinal cord region), which is likely due to the virus being taken up by axons of passage projecting contralaterally (Fig. 7C, axons labeled in the right ventral funiculus (Rt VF)(light blue arrowhead)). Most of the cell bodies labeled were in the thoracolumbar area (Fig. 7D)(cell bodies counted from 13-30 sections per spinal cord region). Strikingly, we found that axons in the right ventral and lateral funiculi (VF, LF) decreased both rostrally and caudally, suggesting that most of these axons are local projections (Fig. 7E)(axons counted from the right and left LF and VF from one section/spinal cord region) and that few thoracolumbar *Atoh1*-lineage neurons project to the LRt, IO, and cerebellar cortex.

**Figure 7.**
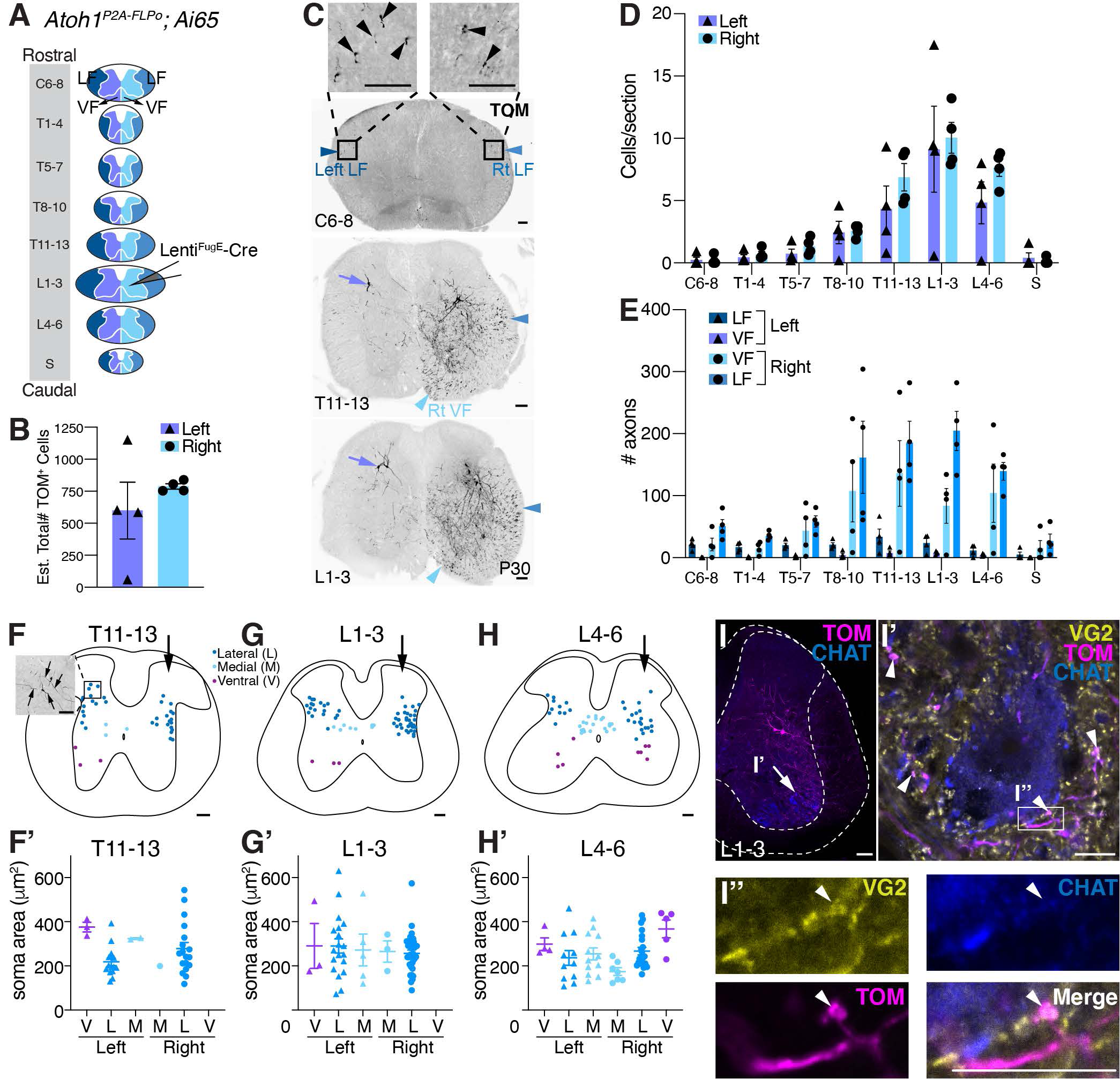
Thoracolumbar *Atoh1*-lineage neurons project locally within the spinal cord. **(A) Diagram** of rostral to caudal sections of the spinal cord (LF (left-dark blue, right-blue); grey matter and VF (left- purple, right-light blue)). Lenti^FugE^-Cre was injected into *Atoh1^P2A-FLPo^;Ai65* mice, such that *Atoh1*-lineage neurons in the right thoracolumbar spinal cord were labeled. (B) Quantitation of the total estimated number of infected cells (TOM^+^) on the left and right sides. The virus appears to be taken up by axons of passage that project from the contralateral side (see C). (C) Representative sections of the spinal cord from a Lenti^FugE^-Cre-injected *Atoh1^P2A-FLPo^;Ai65* mouse. Cell bodies on the right side of the spinal cord and axons in the right (Rt) LF (blue arrowhead) are labeled. Axons in the Rt VF (light blue arrowhead) appear to be axons from cell bodies that are located on the contralateral side of the spinal cord (purple arrows). Some axons in the left LF (dark blue arrowhead) are also seen. (D) L1-3 is the site of peak infection (number of TOM^+^ cell bodies labeled per section) and the number of cell bodies labeled tapers off both rostrally and caudally further away from the injection site. (E) Very few axons on the right side (blue (LF) and light blue (VF)) are detected in rostral sections (C6-8) compared to the site of injection L1- 3. (F-H’) Distribution of infected cells in T11-13, L1-3, and L4-6 regions of the spinal cord (F, G, H, arrows indicate injection on right side). Quantitation of the soma area size of infected medial (M), lateral (L), and ventral (V) *Atoh1*-lineage neurons on the left and right sides of the spinal cord (F’, G’, H’). Inset in F shows two cell bodies labeled on the left side contralateral to the injection that have projections extending dorsolaterally and medioventrally (arrows). (I-I’’) Some of the sparsely labeled thoracolumbar *Atoh1*- lineage neurons have presynaptic terminals near (TOM^+^VG2^+^, arrowheads) or closely apposed (F’’, TOM^+^VG2^+^, arrowhead) to motor neurons (CHAT^+^)(detected in n=4 samples, representative image shown). See Materials and Methods and Results for details of quantitation for B, D, E, F’, G’, H’. Abbrev: LF, lateral funiculus; VF, ventral funiculus. Scale bars: 100 μm (C, I), 10 μm (I’, I’’). Results in graphs presented as mean ± SEM.

Interestingly, the neurons labeled on the left side of the spinal cord, contralateral to the injected side, were located more dorsolaterally than expected for the medial *Atoh1*-lineage population that is known to project contralaterally (Fig. 7C, purple arrows)(Bermingham et al., 2001; Wilson et al., 2008; Yuengert et al., 2015). This observation prompted us to characterize the distribution and cell soma size of infected *Atoh1*-lineage neurons (Fig. 7F-H’). In the spinal cord areas of peak infection (T11-13, L1-3, and L4-6), we found that medial, lateral, and the ill-described ventral *Atoh1*-lineage populations on both sides of the spinal cord were labeled (Fig. 7F-H, n=4, 3-6 sections/n). Quantitation of the soma area found no size differences between the lateral, medial, and ventral population on either side (one-way ANOVA of clusters that had n > 5 cells). We believe that the cells infected on the left side are labeled through their contralaterally-projecting axons that take up the virus on the right side. This suggests that at least a subset of lateral *Atoh1*-lineage cells can also project contralaterally. Indeed, imaging of some of the neurons on the left side shows processes projecting both ventromedially and dorsolaterally (inset in Fig. 7F, arrows), suggesting that these neurons could project both ipsilaterally and contralaterally. While it is possible that cells labeled on the left side are due to diffusion of the virus across the midline, we would have expected the lateral *Atoh1*-lineage neurons in this scenario to project ipsilaterally and therefore, there should be almost equal numbers of axons in the left and right LF given that the number of soma on each side is similar. However, the number of axons in the left LF are low compared to the right LF (Fig. 7E). Therefore, we surmise that the neurons infected on the left side are contralaterally-projecting lateral *Atoh1*-lineage cells, a subset of which could also project ipsilaterally.

Because of the dense ventral projections in our sparsely labeled *Atoh1*-lineage neurons (Fig. 7C, T11-13 and L1-3), we asked whether thoracolumbar *Atoh1*-lineage neurons also synapse on motor neurons. Similar to the cervical *Atoh1*-lineage neurons, we found in all four injections a high density of TOM^+^VG2^+^ puncta (arrowheads, Fig. 7I-I’) near and some very closely apposed to CHAT^+^ motor neurons (arrowhead, Fig. 7I’’). Some of these TOM^+^VG2^+^ puncta might be axo-dendritic synapses (Fig. 7I’, arrowheads), while only a few axo-somatic contacts are detected (Fig. 7I’’). Thus, we find that thoracolumbar *Atoh1*-lineage neurons primarily project locally, where they synapse onto motor neurons, with few neurons projecting to higher brain regions.

### Expression of a silencing allele in caudal Atoh1-lineage neurons leads to a subtle motor defect

To understand how caudal *Atoh1*-lineage neurons affect motor behavior, we expressed GFP- tetanus toxin light chain fusion protein (TeTx), which should inhibit vesicle neurotransmission (Kim et al., 2009), in these neurons using an intersectional genetic strategy (*Atoh1^Cre^;Cdx2::FLPo;R26^LSL-FSF- TeTx^*)(Fig. 8A). In caudal *Atoh1*-lineage neurons heterozygous for the TeTx allele, we found a subtle motor phenotype. We only analyzed mice heterozygous for the TeTx allele because the *Atoh1* gene is close to the *ROSA* locus on chromosome 6 and thus, homologous recombination occurs too infrequently to get homozygosity at the *ROSA* locus along with the *Atoh1^Cre^* allele. Caudal *Atoh1*-lineage neurons expressing the TeTx allele fell off the rotarod sooner compared to controls (Fig. 8B, p=0.0255 for a main effect due to genotype).

**Figure 8.**
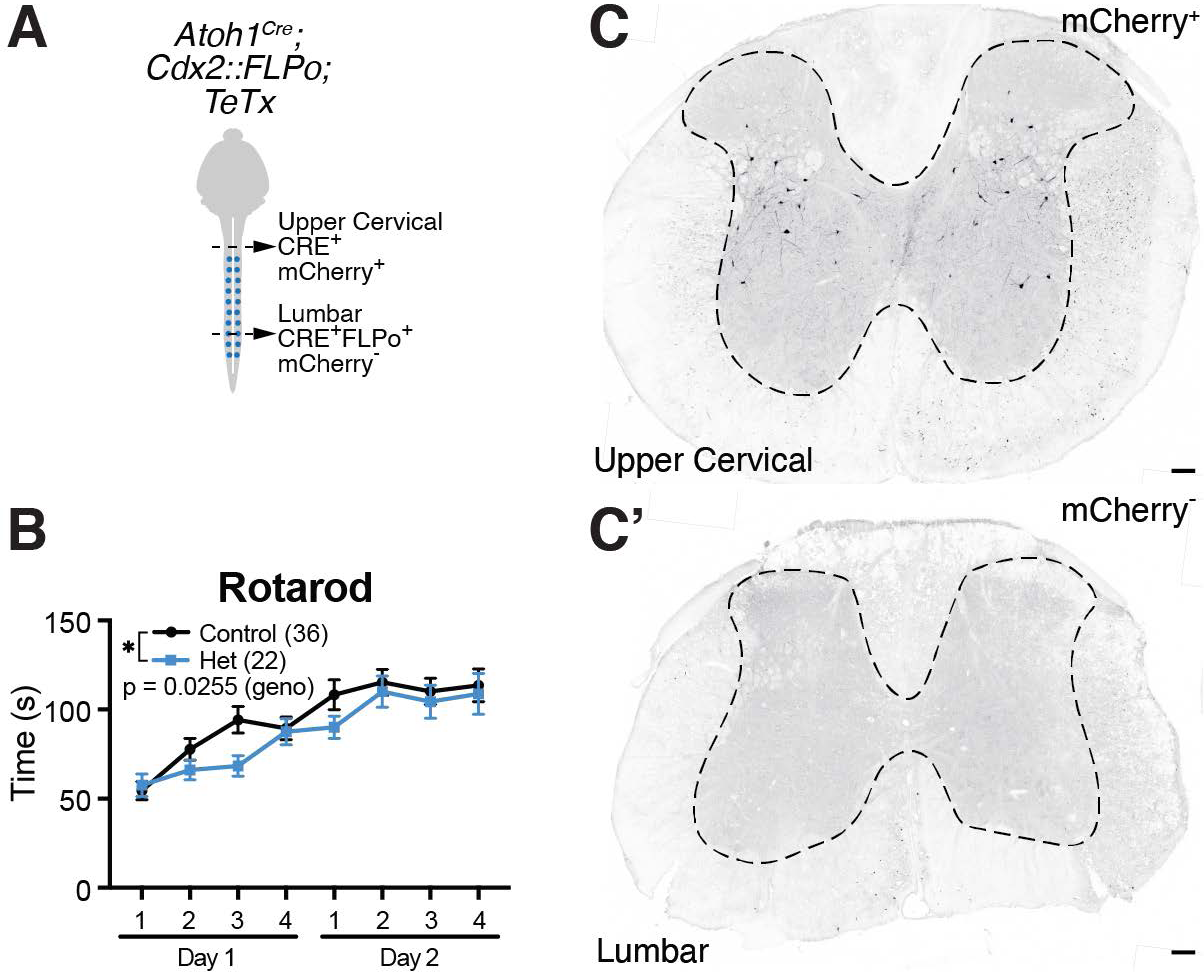
Silencing of caudal *Atoh1*-lineage neurons leads to a subtle motor phenotype. (A) Schematic of dual recombinase strategy to silence caudal *Atoh1*-lineage neurons. (B) Mice heterozygous for the TeTx allele in caudal *Atoh1*-lineage neurons are significantly impaired in the rotarod assay (two- way ANOVA, F(1,448) = 5.025, p=0.0255 for genotype, Control: n=36 (20F:16M), Het: n=22 (10F:12M)). (C-C’) Upper cervical areas of the spinal cord in *Atoh1^Cre^;Cdx2::FLPo;TeTx* mice only have *Atoh1^Cre^* expressed and thus, are mCherry^+^ (C). Lumbar areas of the spinal cord that have both *Atoh1^Cre^* and *Cdx2::FLPo* expressed have lost the mCherry signal (C’). In both panels (C-C’), mCherry signal was amplified using a dsRed antibody. Scale bars: 100 μm (C-C’).

The way that the *R26^LSL-FSF-TeTx^* mouse was designed, CRE recombinase expression allows for mCherry expression while CRE and FLPo recombinase allows for expression of the GFP-TeTx fusion protein (Kim et al., 2009). We performed immunofluorescence and immunohistochemistry with three different GFP antibodies, but were unable to verify expression of the GFP-TeTx fusion protein. Instead, we were able to verify that mCherry was expressed in the upper cervical areas of the spinal cord where *Atoh1^Cre^* is expressed, but not Cdx2*::FLPo* (Fig. 8C-C’). Correspondingly, in the lumbar area of the spinal cord where both *Atoh1^Cre^* and *Cdx2::FLPo* are expressed, the mCherry is no longer expressed indicating that the FLPo recombinase had recombined out the mCherry. All twenty-two heterozygous mice expressing TeTx were confirmed to lack mCherry expression in the lumbar compared to cervical spinal cord.

## Discussion

In this study, we define the anatomy of proprioceptive spinal pathways and find surprising features of these ascending projections (summarized in Fig. 9). We find that CC neurons avoid significant collateralization within the spinal cord, medulla, and CN, although they do collateralize extensively within the cerebellar cortex with some axons crossing the midline. We also discover that cervical *Atoh1*-lineage neurons make up the indirect spino-LRt and spino-olivary pathways while thoracolumbar *Atoh1*-lineage neurons project mostly locally within the spinal cord. In contrast to CC neurons, cervical and thoracolumbar *Atoh1*-lineage neurons make direct connections to MNs and these local connections likely underlie the subtle motor defect seen when a silencing gene is expressed in these neurons. Altogether, we find that the proprioceptive circuits within the spinal cord consist of long-range direct (CC) as well as indirect and locally-projecting (cervical and thoracolumbar *Atoh1*-lineage) neurons that likely mediate different aspects of proprioception.

**Figure 9.**
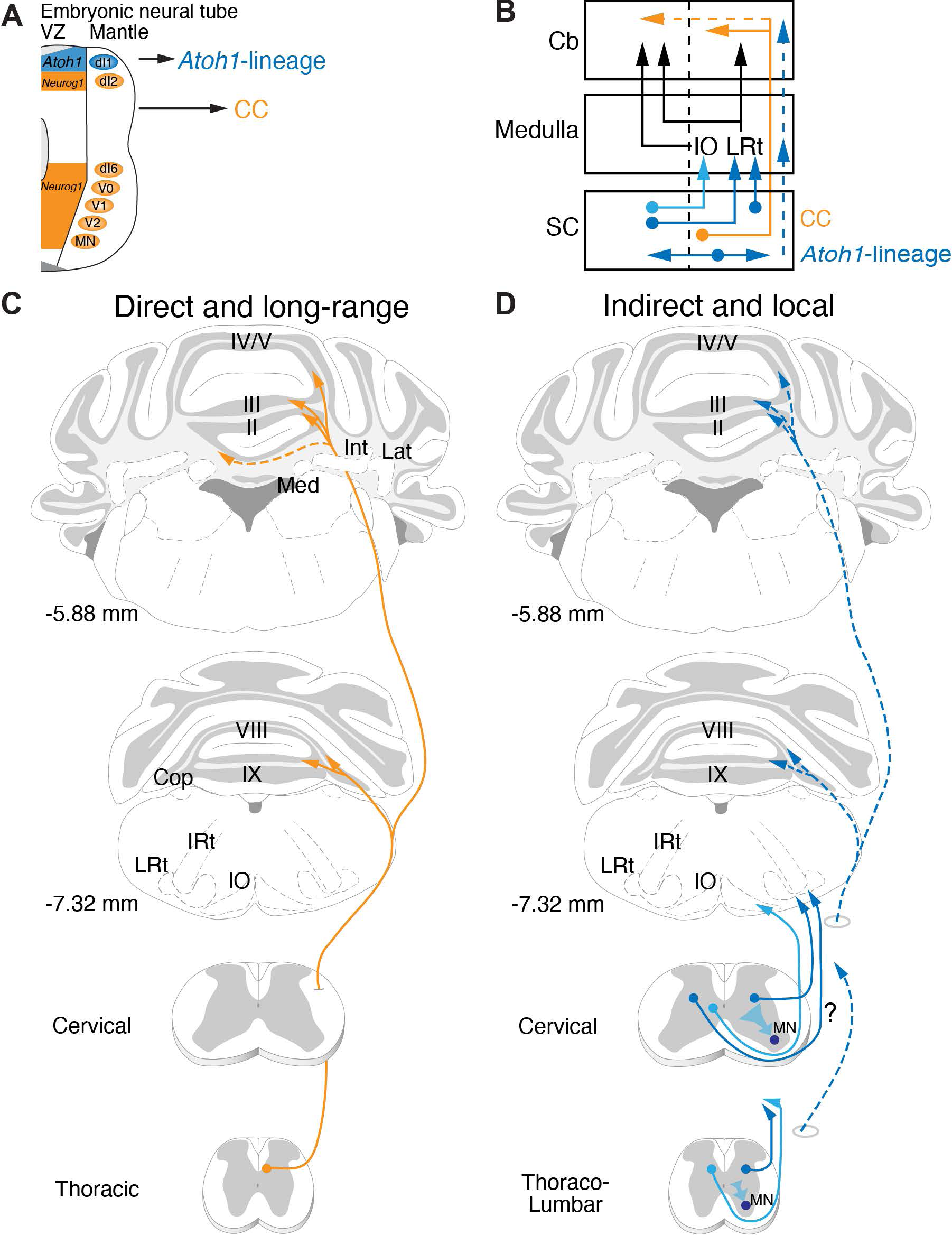
Long-range direct and local indirect proprioceptive pathways. (A) *Neurog1*-lineage neurons of the developing neural tube generate CC neurons that project directly from the spinal cord to the cerebellum while *Atoh1*-lineage neurons form mostly indirect spinocerebellar and local spinal projections. (B) Schematic of major anatomical findings. CC neurons (orange) project ipsilaterally mainly from the thoracic spinal cord directly to the cerebellum. Some CC axons cross the midline (dotted orange). Cervical *Atoh1*-lineage neurons project mainly to the IO and LRt in the medulla (dark and light blue). The LRt neurons then project either ipsilaterally or contralaterally to terminate in the cerebellar cortex as MFs. The neurons in the IO then project contralaterally as well to synapse as climbing fiber (CF) axons onto Purkinje cells (PCs) in the cerebellar cortex. Thoracolumbar *Atoh1*-lineage neurons project mostly within the spinal cord (dark blue, horizontal arrows), although a few project to more rostral regions within the medulla and cerebellum (dotted blue arrows). (C-D) Illustrations of long-range direct and local indirect spinocerebellar pathways. CC neurons project rostrally in the ipsilateral funiculus where they branch extensively in the cerebellum, avoiding the medulla and cerebellar nuclei (Med, Int, Lat) to terminate in vermis I-V, VIII, IXa, and Cop (C). Some CC axons cross the midline within the cerebellum. *Atoh1*-lineage neurons are heterogeneous with diverse projections. Cervical *Atoh1*-lineage neurons in the spinal cord project both ipsilaterally and contralaterally to target mainly the LRt and IO in the hindbrain (D). While we found that some of the *Atoh1*-lineage spino-LRt axons project ipsilaterally, whether some spino-LRt projections come from the contralateral side was not determined in this study (question mark along axons). The spino-IO tract is likely contralateral based on previous literature. *Atoh1*-lineage neurons in the thoracolumbar area project mostly locally within the spinal cord. A few of the axons from thoracolumbar *Atoh1*-lineage neurons may project to the medulla or cerebellar cortex (dotted blue). Presynaptic terminals from *Atoh1*-lineage neurons are found on motor neurons (MN) in both the cervical and thoracolumbar spinal cord. Abbrev: IRt, intermediate reticular nucleus; LRt, lateral reticular nucleus; IO, inferior olive.

### Diversification of proprioceptive information through CC neurons

We find many unique anatomical features of CC neurons that lend insight into how proprioceptive information through CC neurons is relayed. First, we find that a couple thousand CC neurons make up 43-47% of the direct DSCT and VSCT pathways from the AZ relaying hindlimb proprioceptive information. Second, our work clarifies that CC neurons synapse primarily on GCs and do not collateralize significantly within the spinal cord, medulla, or CN, which has been a matter of debate in the literature (Ekerot and Oscarsson, 1976; Szabo et al., 1990; Jiang et al., 2015; Luo et al., 2018). The fact that CC neurons do not collateralize to other parts of the brainstem or CN, as is seen for other MF terminal sources (Sillitoe et al., 2012; Beitzel et al., 2017), suggests that they are not involved in the integration of inputs with other ascending or descending proprioceptive-motor pathways except at the GC level. Lastly, we do find, though, that within the cerebellar cortex, CC neurons extensively diversify their MF terminals between lobules, within a lobule, and even crossing the cerebellum to the contralateral side. Our estimate of approximately 72 MF terminals for 1 CC soma at a population level is similar to the reported 99 terminals/neuron for CC neurons from single axon reconstructions (Luo et al., 2018). Part of the reason for this expansion of proprioceptive information could be for parallel processing across many domains of the cerebellar cortex.

### Atoh1-lineage spino-LRt and spino-olivary neurons

We initially hypothesized that *Atoh1*-lineage neurons made lamina V-SCT or dhSCT neurons based on their anatomical location and developmental studies that reported *Atoh1*-lineage axons go to the cerebellum (Matsushita and Hosoya, 1979; Bermingham et al., 2001; Sakai et al., 2012; Yuengert et al., 2015). Instead, we found that cervical *Atoh1*-lineage neurons make mainly the indirect spino-LRt and spino-olivary pathways. Thus, the cerebellar axonal projections seen during development either extend to the cerebellum and retract during development or die. However, it is possible that *Atoh1*-lineage neurons make a subset of direct spinocerebellar neurons that project to the posterior zone (PZ – lobules VIII/IXa), which was not assessed in our study. Cervical *Atoh1*-lineage neurons could contribute to any of three possible spino-LRt populations that function in posture (bilateral ventral reflex tract), reaching (propriospinal), or grasping (ipsilateral forelimb tract)(Alstermark and Ekerot, 2013; Jiang et al., 2015). We found that cervical *Atoh1*-lineage neurons appear to target the LRt ipsilaterally and as a population can synapse on MNs as well, therefore, it seems likely that *Atoh1*-lineage neurons are at least propriospinal spino-LRt neurons. For the spino-olivary pathway, we found cervical *Atoh1*-lineage neurons targeted areas of the IO (dfDAO, DAO, cMAO^a^, cMAO^b^) consistent with previously described tracing studies and that retrograde tracing from the IO colocalizes with *Atoh1*-lineage neurons in laminae V-VIII (Swenson and Castro, 1983a, b; Matsushita et al., 1992; Oldenbeuving et al., 1999; Flavell et al., 2014). Overall, though, there are numerous other spino-LRt and spino-olivary neurons in the spinal cord suggesting that there are additional sources that are not from the *Atoh1*-lineage (Azim et al., 2014; Pivetta et al., 2014; Choi et al., 2020).

### Local projections of Atoh1-lineage neurons

We find some interesting features of locally-projecting *Atoh1*-lineage neurons and compare and contrast our findings with those of a previous study using an *Atoh1*-enhancer driving CRE recombinase electroporated into mouse embryos (Kaneyama and Shirasaki, 2018). First, we find that thoracolumbar *Atoh1*-lineage neurons project mostly locally, consistent with the findings by Kaneyama and Shiraskai, who found that crossing dI1 axons are intersegmental with only a few traveling far from the soma. Second, we found that *Atoh1*-lineage neurons in both the cervical and thoracolumbar areas of the mature spinal cord make local circuit connections with MNs as previously described during embryogenesis and thus, are a source of pre-motor neurons (Goetz et al., 2015). Kaneyama and Shiraskai found that dI1 commissural axons made synapses on axial MNs, while we found that *Atoh1*-lineage neurons as a population can target the lateral motor column as well. Lastly, similar to our findings, Kaneyama and Shiraskai also found that some dI1 commissural axons come from a fairly dorsolateral population, rather than a medial population, and that these neurons in the lateral population appear to project both contralaterally and ipsilaterally.

### Comparison of motor behaviors mediated by proprioceptive spinal pathways

Loss of proprioceptive input in animal models leads to defects in limb coordination and lack of adaptation to uneven surfaces (Abelew et al., 2000; Windhorst, 2007; Akay et al., 2014). Differences in circuitry between CC (long-range) and caudal *Atoh1*-lineage neurons (indirect and local) suggests that different aspects of proprioceptive-motor behaviors could be mediated by separable microcircuits. We examined the function of CC and caudal *Atoh1*-lineage neurons by expressing a silencing gene, tetanus toxin light chain, in these neurons. Motor function and motor learning were unimpaired when we attempted to silence CC neurons (data not shown), possibly due to sparse expression of our silencing allele, attenuated activity of the silencing allele, or redundancy or compensation of the proprioceptive- motor system. In contrast, we found a slight motor defect in the accelerating rotarod when we silenced caudal *Atoh1*-lineage neurons, consistent with a potential defect in reflexive behavior or postural adjustments that would occur in local circuits.

To assess the function of cervical *Atoh1*-lineage neurons that contribute to the spino-LRt tract, we assessed the behavior of the silenced caudal *Atoh1*-lineage mice in a pellet reaching task (Azim et al., 2014; Becker et al., 2020). We analyzed the reach in five control mice and seven silenced caudal *Atoh1*-lineage littermates, but we did not see any significant differences in pellet reach trajectories due to the low sample size and potentially attenuated function of the TeTx allele (data not shown).

Cervical *Atoh1*-lineage neurons also contribute to the spino-olivary tract. We are not aware of any studies that have manipulated the activity of spino-olivary neurons. However, mice that had glutamatergic signaling in IO neurons knocked out had dystonia-like features, such as twisting, stiff limbs, and tremor, and were unable to perform on the accelerating rotarod test (White and Sillitoe, 2017). We saw no overt twisting, stiff limbs, or tremor in the genetically silenced caudal *Atoh1*-lineage mice, however, these mice did have a defect in the accelerating rotarod and similarly, mice that had *Atoh1* knocked out caudal to the lower medulla were completely unable to perform the rotarod test (Yuengert et al., 2015), suggesting that the spino-olivary system is important for updating motor outcomes. Altogether, caudal *Atoh1*-lineage neurons are heterogeneous comprising parts of the spino-LRt and spino-olivary tracts as well as projecting locally within the spinal cord. Future experiments uncoupling the function of these different components using more robust ablation or acute silencing strategies will lend further insight into the function of discrete proprioceptive circuits.

### Future directions

Our work and others suggest that the spinocerebellar and spino-LRt systems come from several developmental progenitor domains and that any given developmental progenitor domain (*Atoh1*, for example) contributes to several neuronal types (minimally, spino-LRt and spino-olivary tracts, as well as local spinal neurons). Comparing and contrasting features of neurons with similar anatomical connectivity but generated from different progenitor domains may lend insights into the varied functions mediated by seemingly similar anatomical classes. Conversely, separating out different pathways, such as the spino- LRt and spino-olivary tracts that are generated from a single progenitor domain (*Atoh1*) will be important for determining the separate functions of these pathways.

We found that direct CC neurons do not send significant axon collaterals to the LRt or IO and that information to the LRt and IO are in part coming from *Atoh1*-lineage neurons. Therefore, the MF and CF input coming from the spinal cord into the cerebellar cortex comes from different information streams and are not simply collateral copies of the direct CC pathway. Future work focused on how the direct and indirect information streams from the spinal cord either converge or diverge within the cerebellar cortex and the timing with which this information comes in from the two different information streams will be particularly interesting.

## Acknowledgments

This work was supported by R01MH120131 and R34NS121873 to K.M.D., R35HG010719 and R21GM129559 to C.B.G., and the Rita Allen Foundation, Welch Foundation I-1999- 20190330, Kent Waldrep Foundation, R21NS099808 and R01NS100741 to H.C.L. We thank Lin Gan for the *Atoh1^Cre/+^* knock-in mouse, Martyn Goulding for the *Cdx2::FLPo* mouse, Susan Dymecki for the *R26^LSL-FSF-TeTx^* mouse, Mark Behlke and Sarah Jacobi from Integrated DNA Technologies for providing pre-production megamers, Rebecca Seal for the *Vglut1* ISH probe, Thomas Jessell for the *Gdnf* ISH probe, Heankel Cantu Oliveros and Wei Xu for the Lenti^FugE^-Cre virus, Christine Ochoa and Jun Chul Kim for technical assistance, the Neuroscience Microscopy Facility which is supported by the UTSW Neuroscience Dept. and the UTSW Peter O’Donnell, Jr. Brain Institute, LifeCanvas Technologies for tissue clearing assistance, and Jane Johnson, Peter Tsai, Ariel Levine, Euiseok Kim, Abigail Person, and the Lai Lab for helpful discussions and careful reading of the manuscript.

## Multimedia

**Movie 1. *Gdnf^Tom^* cleared spinal cord.** CC cell bodies line the thoracic midline with axons projecting to the lateral funiculus (LF) and then turning rostrally. The *Gdnf^Tom^* mouse line also labels smooth muscle cells lining blood vessels within the spinal cord and along the meninges. TOM^+^ cells along the central canal are also labeled.

**Movie 2. *Gdnf^CreER^; Cdx2::FLPo; Ai65* cleared hindbrain.** CC axons terminate as mossy fibers (MF) in the vermis of lobules I-V, VIII, IXa, and copula (cop). Axons from the spinal cord project directly to the cerebellar cortex avoiding the medulla and cerebellar nuclei.

**Movie 3. *Atoh1^Cre^; Cdx2::FLPo; Ai65* cleared spinal cord.** *Atoh1*-lineage neurons in the spinal cord cluster into medial (M), lateral (L), and ventral (V) populations whose axons travel in the lateral and ventral funiculi (LF and VF). tdTomato is misexpressed in sensory neurons.

**Movie 4. *Atoh1^Cre^; Cdx2::FLPo; Ai65* cleared brain.** The most prominent axonal projections in *Atoh1^Cre^; Cdx2::FLPo; Ai65* mice are to the lateral reticular nucleus (LRt) and inferior olive (IO). Although MF projections are seen in cryosections, they are not obvious in the cleared spinal cord. There is diffuse tdTomato fluorescence in the thalamus and cortex. The misexpression of tdTomato in sensory neurons seen in the spinal cord is seen as projections to the cuneate (Cu), gracile (Gr), and external cuneate nucleus (ECu).

**Movie 5. Three-dimensional projection of cells in the LRt of *Atoh1^Cre^; Cdx2::FLPo; Ai65* mice.** Cropped cell in Fig. 5I is taken from this z-stack of 0.5 μm optical slices with 0.25 μm step. Dense terminations are seen in the LRt. FG (yellow), *Atoh1^Cre^; Cdx2::FLPo; Ai65* axons (TOM^+^, magenta), VG2 (cyan).

**Movie 6. Three-dimensional projection of cells in the IO of *Atoh1^Cre^; Cdx2::FLPo; Ai65* mice.** Cropped cell in Fig. 5J is taken from this z-stack of 0.5 μm optical slices with 0.25 μm step. Dense terminations are seen in the IO. FG (yellow), *Atoh1^Cre^; Cdx2::FLPo; Ai65* axons (TOM^+^, magenta), VG2 (cyan).

